# The molecular genetic landscape of human brain size variation

**DOI:** 10.1101/2022.11.03.514996

**Authors:** Jakob Seidlitz, Travis T. Mallard, Jacob W. Vogel, Younga H. Lee, Varun Warrier, Gareth Ball, Oskar Hansson, Leanna M. Hernandez, Ayan S. Mandal, Konrad Wagstyl, Michael V. Lombardo, Eric Courchesne, Joseph T. Glessner, Theodore D. Satterthwaite, Richard A.I. Bethlehem, Lifespan Brain Chart Consortium, Shinya Tasaki, Bernard Ng, Chris Gaiteri, Jordan W. Smoller, Tian Ge, Michael J. Gandal, Aaron F. Alexander-Bloch

**Affiliations:** Lifespan Brain Institute, The Children’s Hospital of Philadelphia and Penn Medicine, Philadelphia, PA, USA; Department of Child and Adolescent Psychiatry and Behavioral Science, The Children’s Hospital of Philadelphia, Philadelphia, PA, USA; Department of Psychiatry, University of Pennsylvania, Philadelphia, PA, USA; Psychiatric and Neurodevelopmental Genetics Unit, Center for Genomic Medicine, Massachusetts General Hospital, Boston, MA, USA; Stanley Center for Psychiatric Research, Broad Institute of MIT and Harvard, Boston, MA, USA; Department of Psychiatry, Harvard Medical School, Boston, MA, USA; Lifespan Informatics and Neuroimaging Center, University of Pennsylvania School of Medicine, Philadelphia, PA, USA; Department of Psychiatry, University of Cambridge, Cambridge, UK; Department of Psychology, University of Cambridge, Cambridge, UK; Developmental Imaging, Murdoch Children’s Research Institute, Melbourne, Victoria, Australia; Department of Paediatrics, University of Melbourne, Melbourne, Victoria, Australia; Clinical Memory Research Unit, Department of Clinical Sciences Malmö, Lund University, Malmö, Sweden; Memory Clinic, Skåne University Hospital, Malmö, Sweden; Semel Institute for Neuroscience and Human Behavior, University of California Los Angeles, Los Angeles, CA, USA; Wellcome Centre for Human Neuroimaging, University College London, London, UK; Laboratory for Autism and Neurodevelopmental Disorders, Center for Neuroscience and Cognitive Systems @UniTn, Istituto Italiano di Tecnologia, Rovereto, Italy; Department of Neuroscience, University of California, San Diego, San Diego, CA, USA; Autism Center of Excellence, University of California, San Diego, San Diego, CA, USA; The Center for Applied Genomics, The Children’s Hospital of Philadelphia, Philadelphia, PA, USA; Department of Pediatrics, University of Pennsylvania Perelman School of Medicine, Philadelphia, PA, USA; Rush Alzheimer’s Disease Center, Rush University Medical Center, Chicago, IL, USA; Center for Precision Psychiatry, Massachusetts General Hospital, Boston, MA, USA; Department of Genetics, University of Pennsylvania, Philadelphia, PA, USA

## Abstract

Human brain size increases dynamically through early development, peaks in adolescence, and varies up to two-fold among adults. Although previous studies have elucidated changes in brain size across evolution, development, traits, and diseases, the molecular underpinnings of interindividual variation in brain size remain unknown. Here, we leverage postmortem brain RNA sequencing and estimates of brain weight (BW) in 2,531 individuals across three independent datasets, to identify 928 genes that show genome-wide significant associations with either higher or lower BW (BW+, BW-, respectively). These BW gene sets showed distinct neurodevelopmental trajectories and spatial patterns that mapped onto developmental, functional and cellular axes of brain organization. Expression differences among evolutionarily conserved BW genes were predictive of interspecies differences in brain size, and functional annotation of BW genes revealed enrichment for neurogenesis and cell-cell communication. Genome-wide, transcriptome-wide, and phenome-wide association analyses of *in vivo* neuroimaging phenotypes confirmed that the genetic regulation of BW-transcripts influences cortical surface area and volume, as well as behavioral traits related to brain function and disease. Cumulatively, our study represents a major step towards the goal of delineating the causal mechanisms of human brain size variation in health and disease.

## Introduction

The size of the cerebral cortex varies approximately 100-fold across primate species ^1,2^ and 2-fold across adult humans ^3^. While metrics of total brain size have historically been assessed postmortem using tissue weight or volume, highly correlated measurements including volume and surface area are readily quantifiable with modern non-invasive neuroimaging methods. Distinct from head circumference, which plateaus around 5-7 years, total brain volume peaks during adolescence at around 13-15 years. This growth in absolute brain size is accompanied by an increase in the variability of brain size among humans that also peaks in adolescence, and arises from differential contributions of underlying tissue types with discrete growth trajectories ^4^. Many studies have demonstrated high twin-based and single nucleotide polymorphism (SNP) heritability of brain morphology, with many global brain size indices reaching above 50% (twin) and 25% (SNP) of interindividual variance explained by genetic factors ^4–7^. Brain size has also been implicated in myriad clinically relevant contexts, including case-control differences in both neuropsychiatric and neurodegenerative diseases, as well as dimensional associations with anthropometric and cognitive traits ^4,8^.

Large-scale neuroimaging-genetic studies have begun to determine the polygenic architecture of specific components of human brain size, such as total volume and cortical surface area, which are highly genetically correlated ^6,7,9,10^. Collectively, these studies have implicated genes that are involved in critical developmental signaling pathways (e.g., Wnt, PI3K-AKT), are highly expressed during the prenatal period, and are related to neuropsychiatric and neurodegenerative disorders as well as polygenic traits (e.g., cognition). In addition to neuroimaging-genetics, progress towards identifying candidate gene sets and related molecular processes underlying brain size has stemmed from convergent evidence across a diverse body of work, including transcriptome-wide association studies (TWAS) of neuroimaging phenotypes in humans ^11^, blood transcriptomic signatures of neuroimaging-derived brain size metrics in humans ^12,13^, and *in vitro* brain size-related effects in rare neurogenetic and cephalic disorders ^14–16^. In parallel, recent postmortem brain RNA sequencing data-aggregation efforts across tissue banks, such as the PsychENCODE (PEC) project ^17^, have led to the discovery of major transcriptomic signatures of brain evolution and development ^18,19^. Although not direct examinations of brain size, the phylogenetic and ontogenetic dynamics investigated by these studies are intertwined with pronounced changes in brain size across primate species and developmental epochs. In light of these informative prior results, we aimed to directly investigate the functional molecular correlates of interindividual variation in human brain size.

The current study capitalizes on the fact that a direct and accurate measure of brain size – *ex vivo* brain weight – is commonly quantified during postmortem autopsy across tissue banks. We leverage reported brain weight and associated transcriptomic data to perform a genome-wide differential expression analysis of brain weight, in a sample of 2,531 individuals across multiple widely-accessed postmortem brain transcriptomic datasets (PEC discovery N=1,670, GTEx and ROSMAP replication total N=861). We discover replicable associations across independent datasets, identifying 928 genes reaching genome-wide significance that are up-regulated in individuals with higher brain weight (brain weight positive or BW+; N=442 genes) or up-regulated in individuals with lower brain weight (brain weight negative or BW-; N=486 genes). These distinct gene sets show spatiotemporal divergence and relative expression differences across non-human primates, cell-types, and neuropsychiatric disease. Moreover, using an integrative multiscale approach, we pinpoint genetic regulatory mechanisms that link transcriptomic signatures to phenotypic variation in both neuroimaging-derived and brain-related traits. Our comprehensive study represents a significant advance in elucidating the molecular landscape underlying variability in human brain size.

## Results

### Brain size is associated with differential brain gene expression across datasets

We first confirmed the validity of postmortem brain weight as a proxy for brain size. Using comprehensive lifespan neuroimaging models of total cerebrum volume from over 100,000 individuals ^4^ and previous measurements of average brain density ^20^, we found a robust relationship for predicted versus measured brain weight aggregated across datasets (N=3,689, Pearson’s r = 0.92, P < 0.0001; **Fig. S1**; Methods). We saw an almost identical relationship using total cortical surface area in lieu of total cerebrum volume to predict brain weight measurements (r = 0.92, P < 0.0001). In line with many previous studies on brain morphology, we also observed a significant difference in brain weight between sexes across datasets (males > females, t = 25.772, Cohen’s d = 0.76, P < 0.0001).

Next, we performed a transcriptome-wide differential expression analysis of interindividual variation in human brain weight (Methods; **Fig. S2**). Multiple linear regression was used to model the relationship between brain weight and gene expression (N=25,774 genes) from RNA sequencing in postmortem brain tissue (primarily sampled from the frontal cortex; Methods). In the discovery dataset (PEC, N=1,670 subjects), 928 genes showed significant genome-wide differential expression (P_Bonferroni_ < 0.05, i.e., 0.05/25,774, P < 1.94e^-6^; partial r range = -0.16-0.18, absolute partial r range = 0.05-0.18; **Fig. 1A**; **Fig. S3**; **Table S1**). Of these genes, 442 exhibited a positive association with brain weight (i.e., gene expression tended to be greater in individuals with higher brain weights), while 486 genes exhibited a negative association with brain weight (i.e., gene expression tended to be greater in individuals with lower brain weights). These prioritized genes are henceforth referred to as BW+ and BW-genes, respectively. The large majority of both BW+ (86%) and BW- (96%) gene sets had higher than average expression across individuals (**Fig. S4**), suggesting that the differential expression reflected by the BW+ and BW-gene sets can be interpreted as opposing effects of actively expressed genes.

**Figure 1.**
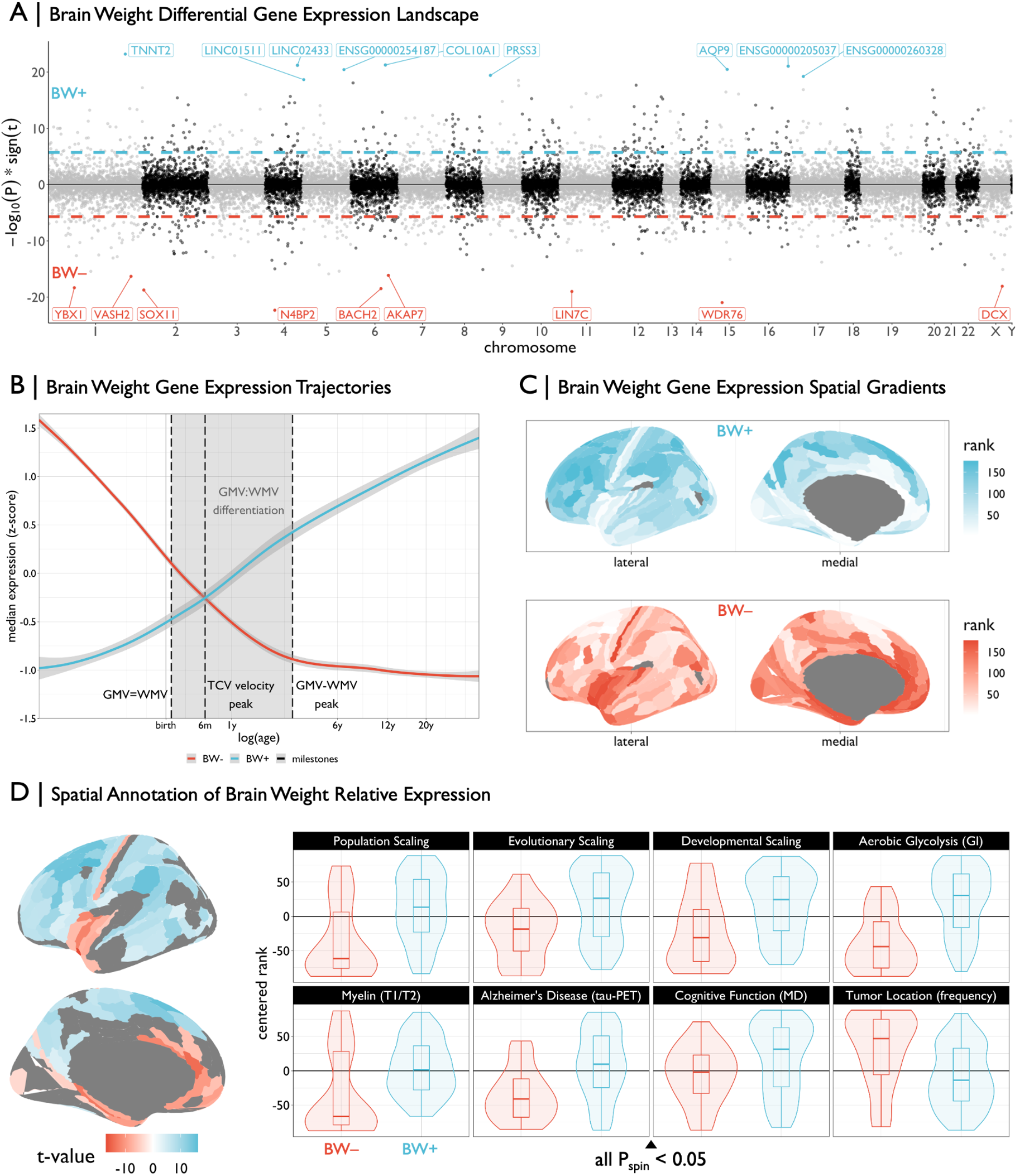
Differential expression of genes associated with an increase or decrease in brain weight across PsychENCODE subjects. **A** | Miami plot showing the relationship between brain weight and expression across all genes (depicting -log10 scaled P-values multiplied by the sign of the t-value effect), ordered by chromosome. Dashed lines represent the genome-wide significance threshold (P_Bonferroni_ < 0.05). For visualization, the top ten genes are labeled for both brain weight positive (BW+, blue) and brain weight negative (BW-, red) sets. **B** | Trajectories of median expression of BW- and BW+ gene sets across development, using the BrainSpan atlas. Black vertical dashed lines represent neuroimaging-derived milestones, highlighting the critical period of gray matter volume (GMV) and white matter volume (WMV) differentiation (shaded area). This period delineates the time when GMV and WMV are equal until the peak difference between GMV and WMV. The brain weight associated gene sets show opposing developmental trajectories, intersecting when the rate of total cerebrum volume (TCV) growth peaks, around 6 postnatal months. Shaded area around trajectories denotes 95% confidence intervals. **C** | Plots of the brain surface showing the differential spatial patterns of the suprathreshold genes (from A) in the Allen Human Brain Atlas (AHBA). Colors denote the rank (from lowest to highest) of cortical regions in terms of median expression of genome-wide significant genes. AHBA probes were resampled from native coordinates to a cortical parcellation (Methods). **D** | (Left) Cortical map of regions showing significant brain weight positive (BW+) versus brain weight negative (BW-) gene expression (P_Bonferroni_ < 0.05) in the Allen Human Brain Atlas (AHBA), i.e., the statistical comparison of the maps in B. (Right) Box-violin plots showing the distributions of diverse multimodal neuroimaging maps across the significant BW+ and BW-regions (see Methods and Fig. S6). For all scaling maps, local/regional surface area was modeled as a function of total cortical surface area across species (evolution), human development, and human subjects (population). All box-violin plots show median and interquartile range (IQR) with whiskers denoting 1.5 × IQR.

We performed multiple sensitivity analyses across subsets of the PEC dataset, showing highly convergent results (**Fig. S4**). For example, when including principal component scores based on genotype (to account for possible ancestry effects) in the BW gene models in the PEC sample, 619 of the BW genes remained significant (P_Bonferroni_ < 0.05), with a high correlation to original model coefficients (r = 0.92, P < 0.0001; **Fig. S4**). There are known inter-relationships between brain size and anthropometrics ^21^, which we observed in our data between brain weight and height, as well as between brain weight and body weight (N=1,206; partial R^2^ from regression model = 0.09 and 0.03, respectively; both P < 0.0001). Consistent with these significant but relatively low effect sizes, BW models including height and weight as covariates (available only in a subset of individuals) were highly correlated to the original models (cross-gene correlation, r = 0.89, P < 0.0001; **Fig. S4**). Comparing genes with significant associations (P_Bonferroni_ < 0.05) across BW and anthropometrics supported the existence of specific associations with BW – 507 of 928 genes were not significant for height or weight (**Fig. S4**). A post-hoc investigation revealed an independent set of genes that showed significant BW-by-age interactions, for which the relationship between BW and gene expression was dependent on age (**Fig. S5**; see Supplemental Information).

Furthermore, a meta-analytic approach combining the two replication datasets (GTEx: N=227 subjects, N=1,327 samples; ROSMAP: N=634 subjects and samples; 733/928 significant BW genes available in all datasets) revealed highly similar BW gene sets: 329/733 PEC BW genes were nominally significant (uncorrected P < 0.05; weighted Sime’s method; ^22^), compared to approximately 37 genes that would be expected to replicate by chance under the null hypothesis of no association ^9^. In addition, combining the discovery and replication datasets revealed highly similar BW gene sets to those identified by the discovery dataset alone (weighted Sime’s method: 733/733 overlapping genes, P_Bonferroni_ < 0.05), with an additional 116 genes meeting significance under the combined framework (P_Bonferroni_ < 0.05; **Table S2**).

### Brain size associated genes have distinctive neurodevelopmental and anatomical profiles

Comparing BW+ and BW-genes revealed opposing neurodevelopmental trajectories in tissue samples spanning the prenatal period through young adulthood in the BrainSpan atlas (**Fig. 1B**; ^23^). Moreover, the BW sets showed striking overlap with previously defined gene sets based on developmental trajectory across the perinatal period in an independent dataset with dense prenatal sampling (BW+ and “rising” genes, BW- and “falling” genes; **Fig. S5**; ^24^). For example, *OPALIN* (BW+ gene) was the highest-ranked “rising” gene based on the slope of expression, and *SOX11* (BW-gene) was the 4th highest-ranked “falling” gene. The fact that neither BW gene set was relatively highly expressed just after birth, at the peak of overall brain growth and during a critical period of constituent tissue class differentiation (**Fig. 1B**), is consistent with previously established findings of low overall gene expression variation after the late-fetal transition ^18^.

In line with the neurodevelopmental results, BW+ and BW-gene sets revealed opposing spatial topographies of average gene expression in densely sampled postmortem adult human brain tissue (**Fig. 1C**; ^25^), with BW+ genes showing high expression in association cortical regions and BW-genes showing high expression in sensorimotor and paralimbic/limbic cortical regions. Therefore we computed a map of relative expression differences, comparing BW+ versus BW-genes across cortical regions. Out of 177 cortical regions with sufficient samples (Methods), 115 (65%) showed significantly greater relative expression of either BW+ or BW-gene sets (two-tailed t-test per region, P_Bonferroni_ < 0.05 across regions). Out of 12 subcortical regions, 10 showed significant BW-relative expression differences (**Table S3**). These differences in BW +/- relative expression demonstrated extreme regional divergence and were far greater than expected by chance given the number of BW+ and BW-genes (P_permutation_ < 0.0001). Moreover, these regions of significant BW +/- relative expression (**Fig. 1D**, left) showed different patterns of correspondence with multiple maps of brain organization, function, and disease (all P_spin_ < 0.05; **Fig. 1D**, right) – including axes of fine-grained cortical surface area expansion ^26,27^, aerobic glycolysis ^28^, myelin-related topography ^29^, cognitive function ^30^, tau distribution in Alzheimer’s disease patients ^31^, and tumor location frequency across patients with glioblastoma and low grade glioma ^32^ (**Fig. S6**). Additionally, the BW +/- relative expression map was robust to an alternative cortical parcellation (**Fig. S5**), and regions with high relative expression of BW genes showed greater morphological effects across diagnostic categories and showed differential patterns of neurodevelopmental growth. Specifically, the spatial topography of BW +/- relative expression was correlated with regional surface area alterations across neuropsychiatric disorders (r = 0.52, P = 0.0017, P_permutation_ < 0.0001, P_spin_ < 0.0001; **Fig. S6**; ^33^); as well as regional variation in the age at peak gray matter volume during neurodevelopment (r = -0.47, P = 0.0046, P_permutation_ < 0.0001, P_spin_ < 0.0001; **Fig. S6**; ^4^).

Cortical regions with BW +/- relative expression differences were also affiliated with specific neurophysiological, functional and cytoarchitectonic classes. Using available spatially-comprehensive positron emission tomography (PET) neuroradiography data ^34^, areas of high relative expression of BW-genes showed greater densities of dopaminergic and serotonergic receptors compared to areas of high BW+ relative expression (**Fig. S7**). Moreover, there was an abundance of BW-regions in primary sensory cortices containing short distance (<80mm) white matter fibers as estimated with diffusion MRI, and an abundance of BW+ regions in higher-order association cortices with long distance white matter fibers (**Fig. S7-8**). Relatedly, a 3D-reconstructed Merker-stained postmortem human brain ^35^ revealed a highly diverging laminar architecture of relative cell density, but not relative thickness, in layer IV for BW-regions and layers V/VI for BW+ regions ^36^, a pattern which was also validated using spatial RNA sequencing data (**Fig. S9**; ^37^). Collectively, these results are aligned with century-old observations on cytoarchitectonic variation across the cerebral cortex ^38^, demonstrating the convergence of BW-regions with the relatively small number of regions in “heterotypic” cortex and BW+ regions with the more abundant canonical “homotypic” 6-layered cortical regions (**Fig. S9**).

### Brain size associated genes show cross-species and cell-type specificity

Genes that are positively associated with variability in brain size among humans may also be implicated in brain size on an evolutionary timescale. Using data from a parallel cross-species study in humans and non-human primates (http://evolution.psychencode.org/; N=16 regions; N=6 humans [H], not included in main analyses; N=5 chimpanzees [C]; N=5 macaques [M]; ^19^), we sought to determine whether phylogenetic variability in brain size (PBS+ = H > C > M; PBS-= H < C < M) was associated with differential expression in BW genes. Across brain regions, 726 genes showed patterns of interspecies differential expression differences in accordance with PBS, and over 60% of genes previously identified as differentially expressed in humans ^39^ showed specific PBS directional effects (Methods). We found a general consistency between overlapping BW and PBS genes (N=366 significant gene-by-region effects, N=110 unique genes) such that BW+ aligned with PBS+ and BW-aligned with PBS-, respectively (73% congruence, P_permutation_ < 0.0001; **Fig. 2A**). Considering all homologous genes across species, we found that there was a moderate but significant relationship between BW effect size and PBS effect size (Spearman r = 0.19, P_permutation_ < 0.0001). It is interesting to note that in macaques, there were no regions with significant BW+ relative expression (**Table S4**; ^40^), consistent with the hypothesis that BW+ genes are implicated in human-specific cortical expansion.

**Figure 2.**
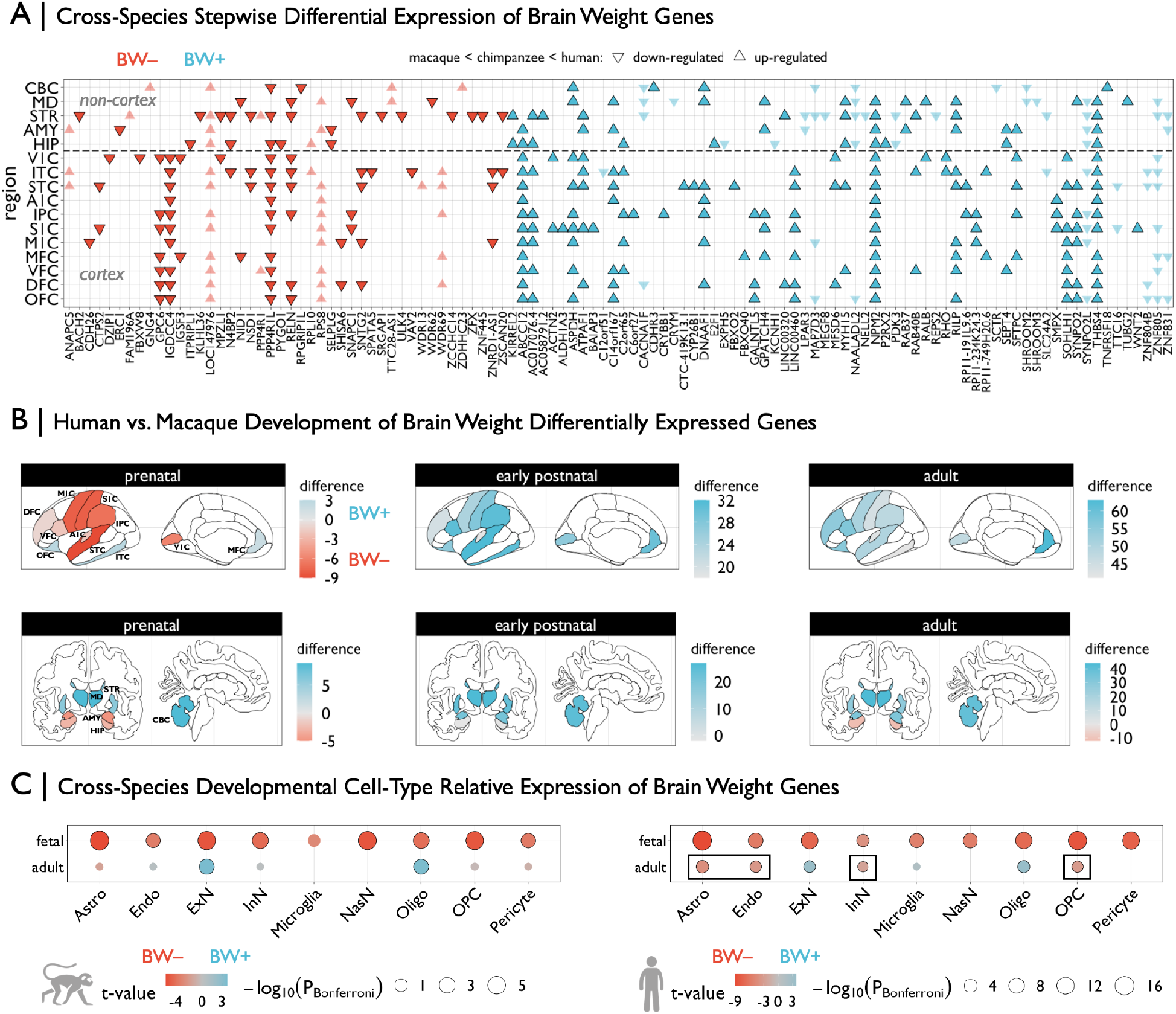
Genes associated with brain weight show differential effects across species. **A** | Grid plot showing brain weight (BW) associated genes that are also significantly differentially-expressed in humans relative to chimpanzees and macaques (P_Bonferroni_ < 0.05), across 16 brain regions (11 neocortical areas). These are BW-associated genes that were previously defined ^39^ as having higher or lower expression compared to the two species collectively, and reanalyzed to assess stepwise interspecies effects which reflect the absolute differences in brain size between species (i.e., up-regulated implies gene expression humans > chimpanzees > macaques, while down-regulated implies gene expression humans < chimpanzees < macaques). Triangles represent directions of effects and colors denote the respective BW gene set. Genes that are highlighted show congruent directional effects in respective BW sets (i.e., BW+ and upregulated in humans, and vice versa). **B** | Brain plots of differences in counts of number of BW+ or BW-genes that were defined to be significantly differentially-expressed in humans relative to macaques each of three developmental epochs – a positive regional difference (blue) indicates that BW+ genes tend to be upregulated in humans in that region in a given developmental epoch, while a negative difference (red) indicates that BW-genes tend to be upregulated. The same 16 regions from A are shown anatomically, based on manual assignment using a common human atlas ^47^. **C** | Differential expression of BW+ versus BW-genes across individual cell-types, using cell-specific RNA sequencing data in fetal and adult samples from macaques (left) and humans (right). BW-relative expression (red) indicates that BW-genes are more highly expressed in that cell type compared to BW+, whereas BW+ relative expression (blue) indicates the opposite effect. Black outlines denote significant effects (P_Bonferroni_ < 0.05). Circles are scaled according to Bonferroni-corrected p-values. Black rectangles denote human-specific effects relative to macaques. V1C: primary visual cortex, M1C: primary motor cortex, S1C: primary somatosensory cortex, A1C: primary auditory cortex, ITC: inferior temporal cortex, IPC: inferior parietal cortex, STC: superior temporal cortex, OFC: orbitofrontal cortex, VFC: ventrolateral frontal cortex, DFC: dorsolateral frontal cortex, MFC: medial frontal cortex, STR: striatum, MD: medial dorsal thalamus, AMY: amygdala, HIP: hippocampus, CBC: cerebellar cortex. Astro: astrocytes, Endo: endothelial cells, ExN: excitatory neurons, InN: inhibitory neurons, NasN: nascent neurons, Oligo: oligodendrocytes, OPC: oligodendrocyte precursor cells.

If the divergent neurodevelopmental trajectories of BW+ and BW-genes are evolutionarily conserved, convergent differential expression patterns in the BW gene sets would be predicted between humans and non-human primates. We leveraged established differential expression statistics between humans and macaques ^19^ to assess whether there were cross-species differences in BW +/- relative expression across brain regions and developmental epochs. Consistent with human developmental patterns (**Fig. 1C**), greater numbers of BW-genes were differentially-expressed across species in the prenatal epoch and greater numbers of BW+ genes were differentially-expressed across species in the postnatal epochs (**Fig. 2B**). Although there were more differentially-expressed BW+ genes in the early postnatal period in humans (**Fig. S10A**), there were a small number of exceptions to the general pattern observed in the prenatal period (generally BW-predominant) and the adult period (generally BW+ predominant). Specifically, the striatum and thalamus showed more BW+ differentially-expressed genes prenatally, and the hippocampus showed more BW-differentially-expressed genes in adult humans. This relative heterochronicity in the abundance of BW differentially-expressed genes in non-cortical brain regions could reflect the protracted neurogenesis of adult hippocampal compared to thalamic neurons ^41^.

We hypothesized that BW +/- relative expression differences could be related to the relative abundance of specific cell types among brain regions and across developmental and evolutionary timescale. In single cell (fetal) and single nucleus (adult) RNA sequencing data ^18^ in macaques (http://evolution.psychencode.org/) and humans (http://development.psychencode.org/), there was greater BW-relative expression across most macaque prenatal cell types and all human prenatal cell types, including human-specific progenitor cell subtypes (**Fig. 2C**; **Fig. S10B**). Overall, in both species, there were fewer adult cell types with BW +/- relative expression differences. Whereas BW-genes did not show any relative expression differences in adult macaques, they demonstrated greater relative expression in human endothelial and oligodendrocyte progenitor cells, as in their respective prenatal cell type counterparts. In both species, BW+ genes showed greatest relative expression in excitatory neurons (primarily supragranular layer subtypes) and oligodendrocytes, in contrast to BW-relative expression in those prenatal counterparts (**Fig. 2C**; **Fig. S10B**). These results are consistent with spatiotemporal and laminar findings (**Fig. 1**; **Fig. S6**), as well as the differential trajectories of human gene expression across BW+ (low prenatal, high postnatal) and BW- (high prenatal, low postnatal) gene sets. Collectively, this coincides with observed protracted maturation of cortical myelin ^4^ as well as the relative neuronal density of cortical regions with expanded supragranular layers ^38^.

The BW cell-type divergence is further reflected by the predominance of *prenatal* BW-differentially-expressed cell types *within* species versus a predominance of *postnatal* BW+ differentially-expressed genes *across* species. The human-specific neotenic BW-differential expression in inhibitory neurons and astroglial cell types is in line with the abovementioned findings of BW-relative expression between species in the adult hippocampus – a structure which not only is preferentially enlarged in humans relative to non-human primates ^42^ but is also highly connected (via specified GABAergic inhibitory circuits ^43^) to functionally related areas of significant BW-relative expression (see **Fig. 1D**). Independent human fetal RNA sequencing across brain regions further suggest greater BW-relative expression signatures in the subventricular zone (**Fig. S11**), which is expanded in primates ^44,45^, and in line with the BW-enrichment of neuronal precursor subtypes and perseveration of BW-enrichment in adult human inhibitory neuron subtypes ^46^.

### Brain size associated genes are implicated in neuropsychiatric and neurodevelopmental disorders

Given myriad previous studies showing brain volume abnormalities across a range of neuropsychiatric and neurodegenerative disorders, we predicted that BW gene sets would overlap with genes that are differentially expressed in patients. Across shared genes in each condition (N=4,226 genes), there were significant gene-level correlations between PEC BW-expression statistics and differential expression (DE) statistics from previous studies on neuropsychiatric ^48^ and neurodegenerative ^49,50^ disorders (**Fig. 3A**). Specifically, positive correlations were found with alcohol abuse disorder (AAD), whereas negative correlations were found with Parkinson’s disease (PD), Alzheimer’s disease (AD), autism spectrum disorder (ASD), bipolar disorder (BD), and schizophrenia (SCZ), with no significant correlation with inflammatory bowel disease (IBD) – a non-neural control. The direction of correlation related to the proportion of intersecting significant BW and DE genes (both thresholded P_FDR_ < 0.05) showing either convergent (positive correlation = BW/DE +/+ or -/-) or divergent (negative correlation = BW/DE +/- or -/+) effects, with five disorders showing significant BW-DE alignment scores (all P_permutation_ < 0.001; **Fig. 3B**). Moreover, we identified a subset of 36 BW-DE genes shared across disorders showing divergent BW-DE alignment (AD, ASD, BD, SCZ; **Fig. 3C**). This subset of BW-DE genes were almost perfectly separated in terms of the direction of effect, such that BW+ genes were downregulated and BW-genes were upregulated in patient tissue (**Fig. 3C**). In supplemental analyses, we also derived a cumulative BW transcriptome score, which showed case-control differences across diagnostic categories (**Fig. S12**). Consistent with the spatial alignment between the map of BW+/- relative expression and tau-PET (**Fig. 1D**), the cumulative transcriptome score was also associated with indices of tau and amyloid neuropathology in patients with dementia in the independent ROSMAP dataset (Supplemental Information).

**Figure 3.**
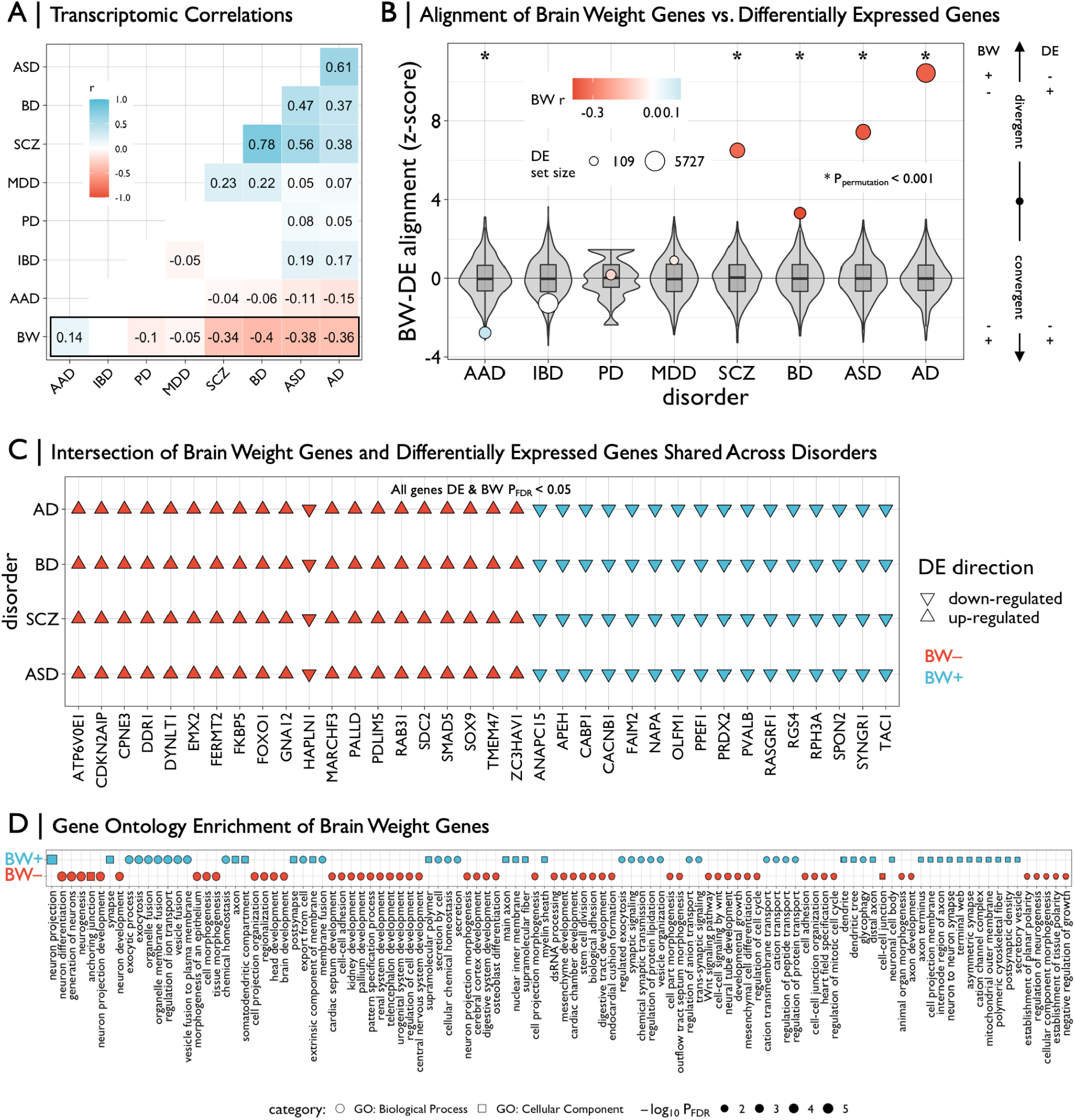
Brain weight gene sets are differentially enriched across disease and functional domains. **A** | Matrix of transcriptomic correlations of differential expression (DE) statistics in multiple human diseases and brain weight (BW) statistics in the PsychENCODE dataset. Pairwise Pearson’s r coefficients were computed across overlapping genes between datasets (N=4,226) – only significant (P_Bonferroni_ < 0.05) values are plotted. Autism spectrum disorder (ASD), bipolar disorder (BD), schizophrenia (SCZ), major depressive disorder (MDD), Parkinson’s disease (PD), alcohol abuse disorder (AAD), Alzheimer’s disease (AD). Inflammatory bowel disease (IBD) was included as a non-neural control. **B** | Plot showing the alignment of BW and DE genes within each disorder. Intersecting lists of significant BW and DE genes (both P_FDR_ < 0.05) were categorized as “convergent” or “divergent” based on the concordance of the directions of effects, then z-scored according to a null distribution based on 10,000 resamples of BW gene sets of similar size (gray box-violins). Circles are sized according to the number of DE genes and colored according to values in A. Asterisks denote significant (P_permutation_ < 0.001) z-scores. **C** | Grid plot showing 36 significant BW and DE genes (DE & BW P_FDR_ < 0.05) in patients with AD, BD, SCZ, and ASD. Triangles represent directions of effects (up- or down-regulated in patients compared to controls) and colors denote the respective BW gene set. **D** | Grid plot showing significant (P_FDR_ < 0.05, reduced for visualization, see **Table S5**) Gene ontology enrichment of brain weight (BW) associated genes for biological processes (circle) and cellular components (square) using ToppGene ^57^. Shapes are sized and ordered (high-to-low) according to adjusted negative log-scaled p-values. All box-violin plots show median and interquartile range (IQR) with whiskers denoting 1.5 × IQR.

Furthermore, functional gene ontology enrichment analysis of the BW gene sets aligned with disease differential expression signatures. BW+ genes showed significant enrichment for axonal and synaptic functions, including ion channel signaling, and BW-genes showed significant enrichment for neurogenesis/organogenesis and abnormal developmental physiology (**Fig. 3D, Fig. S13**; **Table S5**). In addition, BW genes showed enrichment across multiple modules derived from gene co-expression networks in the PEC dataset ^51^ (**Fig. S14A**), indicating that BW genes are significant components of transcriptional regulatory programs, some of which may be cell-type specific. In alignment with the cell-type enrichment results described above, we found module-specific enrichments of BW-genes in “geneM3” (related to astrocytes) and of BW+ genes in “geneM1” and “geneM2” (related to neuronal/synaptic signaling and oligodendrocytes, respectively). Using gene co-expression modules previously derived In the ROSMAP cohort ^52^, we found module-specific enrichments of BW-genes in “m107” (related to astrocytes) and of BW+ genes in “m23” (related to neuronal/synaptic signaling) (**Fig. S14A**; **Table S6**). Moreover, we found enrichment of BW-genes in “m109”, which was shown in prior work ^52^ to be linked to cognitive decline and multiple indices of neuropathology (**Fig. S14B**). Further gene set enrichment analysis using independent RNA-seq data across multiple tissues from the Human Protein Atlas ^53^ yielded significant enrichment of BW+ genes in the adult cerebral cortex versus enrichment of BW-genes in female reproductive tissues (**Fig. S15A**), which aligns with the high prenatal expression of BW-genes.

To examine the landscape of rare genetic variation in BW gene sets, hypothesized to be associated with neurodevelopmental disorders, we leveraged two metrics of mutational constraint derived from public databases of whole exome sequencing and de novo mutations ^54,55^. We found that BW-genes, and not BW+ genes, were significantly enriched for intolerance to loss-of-function (protein truncating) variation (**Fig. S15B**). Using previously defined gene sets of high dosage sensitivity (N=2,987 haploinsufficient; N=1,559 triplosensitive), we found a significant bidirectional enrichment (χ^2^ = 6.1534, P = 0.0131, P_permutation_ = 0.0047; **Fig. S15C**), with greater overlap in BW- and haploinsufficient genes and between BW+ and triplosensitive genes ^56^. Moreover, BW-genes showed greater overlap with documented pathogenic de novo developmental mutations (BW- OR = 1.46, P_permutation_ = 0.008; BW+ OR = 0.83, P_permutation_ = 0.31), which are associated with abnormalities of brain and cognition, head size, facial, gastrointestinal, and reproductive systems (all P_permutation_ < 0.05; **Fig. S15D**).

### Brain size associated transcription is regulated by genetic variants

The genetic regulation of the transcriptome is one factor that could contribute to interindividual variability in brain gene expression and brain size. We used multiple approaches to assess the genetic regulation of BW transcripts. First, leveraging individual-level transcriptome and genotype data from subjects in PEC as the reference dataset, we performed a transcriptome-wide association study (TWAS) of structural neuroimaging phenotypes in UK Biobank (N=21,936), normalized using BrainChart ^4^. For total cerebrum volume (TCV) and cortical surface area (SA), two of the phenotypes whose models accurately predicted brain weight, 21 genes reached genome-wide significance (P_Bonferroni_ < 0.05) with 15 shared between both phenotypes and an additional 6 significant only for SA. 14 of these shared genes were not identified in previous TWAS of related phenotypes – intracranial volume and total brain volume ^11,58^. Of these 15 significant shared genes, 2 genes (*PRR13, EGFR*) overlapped with BW gene sets (OR = 3.07, P_permutation_ = 0.0053). Sensitivity analysis (conducted with an uncorrected P < 0.05 for TWAS genes) also demonstrated significant enrichment of TCV and SA genes, with 40 genes overlapping with BW gene sets (OR = 9.21, P_permutation_ < 0.0001). Supplemental analyses with a wide range of neuroimaging phenotypes including diffusion MRI measures (**Fig. S16**), at both whole-brain (**Table S7**) and regional (**Table S8**) scales, also demonstrated significant overlap with BW genes (P_Bonferroni_ < 0.05; see Supplemental Information; see also **Table S9**).

To further evaluate the genetic influence of BW gene sets on other phenotypes, we leveraged >2,300 GWAS summary statistics across micro- and macro-structural neuroimaging phenotypes ^10^. Using established gene-set enrichment methods (MAGMA; ^59^), competitive enrichment analyses of multimodal global neuroimaging phenotypes (averaged or summed across regions) revealed a strong and specific link of BW-genes to brain size (cortical surface area and volume) but not brain composition measured via diffusion MRI (**Fig. 4A**; **Table S10**). Interestingly, we did not find enrichment of BW gene sets with recent large-scale GWAS of height ^60^, body mass index ^61^, or head size (head circumference and intracranial volume; ^62–64^), which could either indicate insufficient statistical power for current head size GWAS or suggest a specificity to brain parenchyma size (**Table S11**). Based on BW-gene set enrichment in GWAS of morphological phenotypes (cortical surface area and volume), post hoc analysis revealed a spatially patterned enrichment of BW-genes in GWAS of specific cortical regions (**Fig. 4A**; **Table S12**). Remarkably, cortical regions showing significant BW-gene enrichment in surface area GWAS overlapped with regions with high BW-relative expression as shown in Fig. 1D (odds ratio (OR) = 6.17, P = 1.14e-5, P_spin_ < 0.0001), but did not overlap with regions with high BW+ relative expression (OR = 0.61, P = 0.14, P_spin_ = 0.94). These brain regions with significant BW-gene enrichment in surface area GWAS also showed significant overlap with established regions of hypo-allometric scaling (OR = 3.98, P_spin_ < 0.0001; using maps of allometric (non-linear) scaling of regions with total brain size across population, evolutionary, and developmental scales; **Fig. S5**). Collectively, these results integrate neuroimaging maps of local nonlinear scaling with brain size, the genetic regulatory signatures of brain size-related morphology, and the spatial differential gene expression underlying brain size variation.

**Figure 4.**
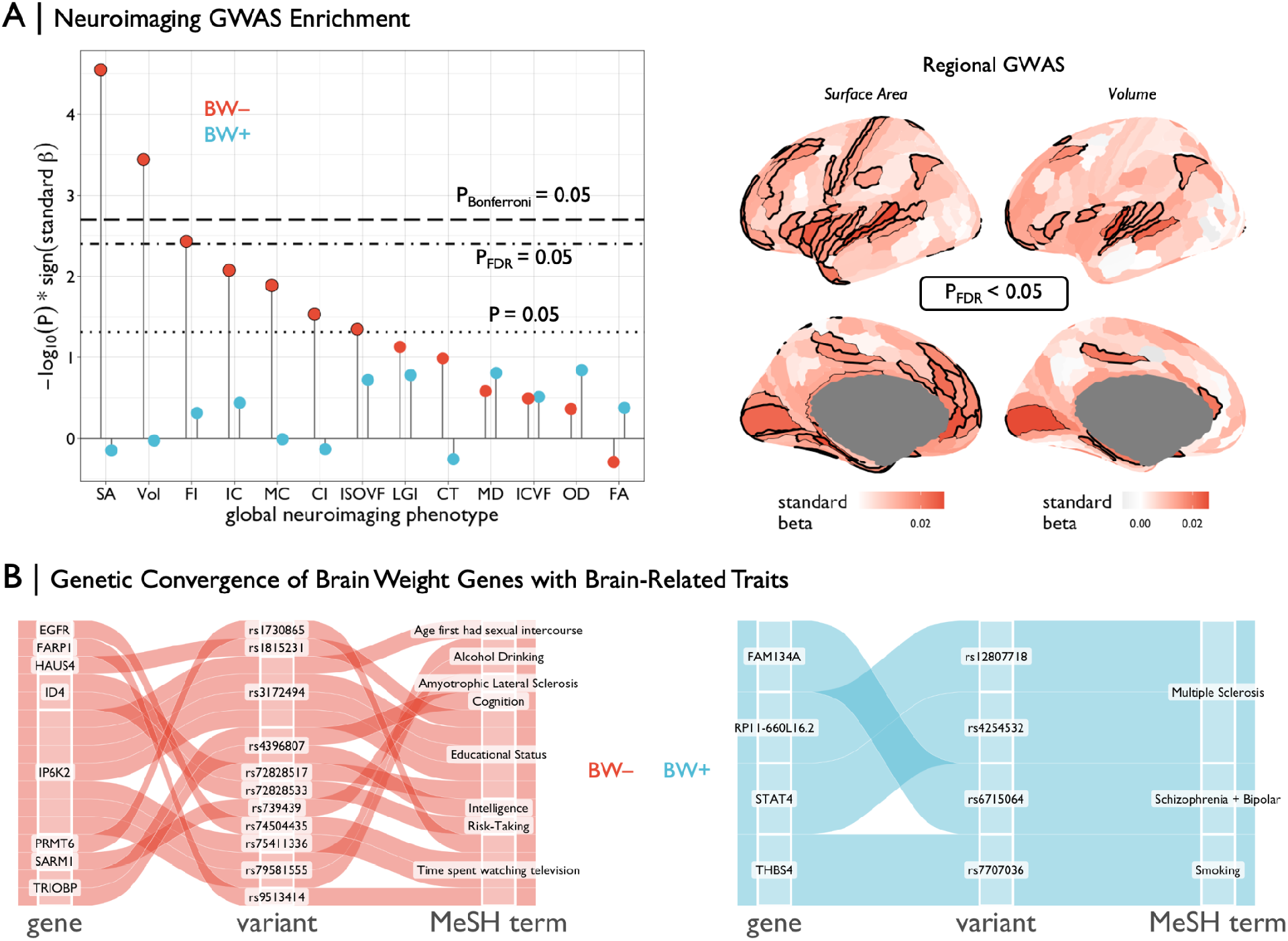
Functional relevance of genes associated with brain weight. **A** | (Left) Lollipop plot showing MAGMA ^59^ enrichment statistics for BW gene sets across Genome-Wide Association Studies (GWAS) of 12 multimodal neuroimaging phenotypes we generated in the UK Biobank dataset (Methods). Black lines represent various significance thresholds. Black outlines denote suprathreshold effects (at least uncorrected P < 0.05). No significant effects were observed for the BW+ gene set. (Right) Enrichment of BW-genes for regional surface area and gray matter volume across 180 cortical regions (the same parcellation as in Fig. 1; Methods). Black outlines denote significant areas of enrichment (P_FDR_ < 0.05 shown for visualization, see Table S10). **B** | Alluvial diagrams representing the concordance between BW genes (P_Bonferroni_ < 0.05) and multivariate multiple quantitative trait loci (mmQTL; ^65^) with genome-wide significant association with brain-related traits. SNP: single nucleotide polymorphism, FDR: false discovery rate.

Finally, we used recently published multivariate multiple quantitative trait loci (mmQTLs) derived from the same three cohorts ^65^ used in the current study to triangulate genetic variants (i.e., single nucleotide polymorphisms) that may influence brain-related traits via an impact on regulating expression of BW genes. Thus, we were able to identify 8 BW-genes (*EGFR, FARP1, HAUS4, ID4, IP6K2, PRMT6, SARM1, TRIOBP*) and 4 BW+ genes (*FAM134A, RP11-660LI6*.*2, STAT4, THBS4*) with mmQTLs associated with brain-related traits including educational attainment, impulsivity and psychotic disorders (**Fig. 4B**). Of these, BW-genes *ID4, IP6K2*, and *TRIOBP* showed evidence of pleiotropy across multiple brain-related traits, and expanded analysis across all studied traits showed additional pleiotropic associations between both BW gene sets and multiple anthropometric and metabolic domains (**Table S13**).

### A brain size transcriptomic score is associated with clinical outcomes in healthcare system data

To further investigate the clinical correlates of genetic variation influencing BW transcription, we conducted a phenome-wide association study (PheWAS) of patient medical outcomes in the Mass General Brigham (MGB) Biobank, an independent biorepository from a U.S. healthcare system ^66^. Inspired by recent work on the integration of transcriptomic data and polygenic indices ^67^, we calculated a cumulative BW-transcriptome score for MGB Biobank patients as a coefficient-weighted sum of imputed gene expression (Methods). We then used logistic regression models to examine relationships between the BW-transcriptome score and 1,482 case-control disease/disorder phenotypes in up to 37,272 individuals of European ancestry. All models included sex assigned at birth, current age, genotyping batch, and the first 10 genetic principal components as covariates.

Overall, we found that 241 phenotypes were associated with the BW-transcriptome score at P_FDR_ < 0.05, of which 36 were P_Bonferroni_ < 0.05, with odds ratios ranging from 0.3 to 3.1 per standard deviation increase in the score (**Fig. 5**; **Table S14**). The most notable associations were with neurodegenerative disorders (negative association), general mental and behavioral problems (negative), congenital brain abnormalities and neural tube defects (negative), and intellectual disability (positive). Individuals with extreme BW-transcriptome scores in both directions also showed worse health in many other bodily systems. They were more likely to suffer, for example, from infertility or adverse pregnancy outcomes. Collectively, this integrative genomic scoring approach, leveraging observed BW-expression associations and imputed gene expression data, demonstrates a proof-of-concept bridge between functional enrichments and bioinformatic annotations of the BW gene sets with real-world translational potential.

**Figure 5.**
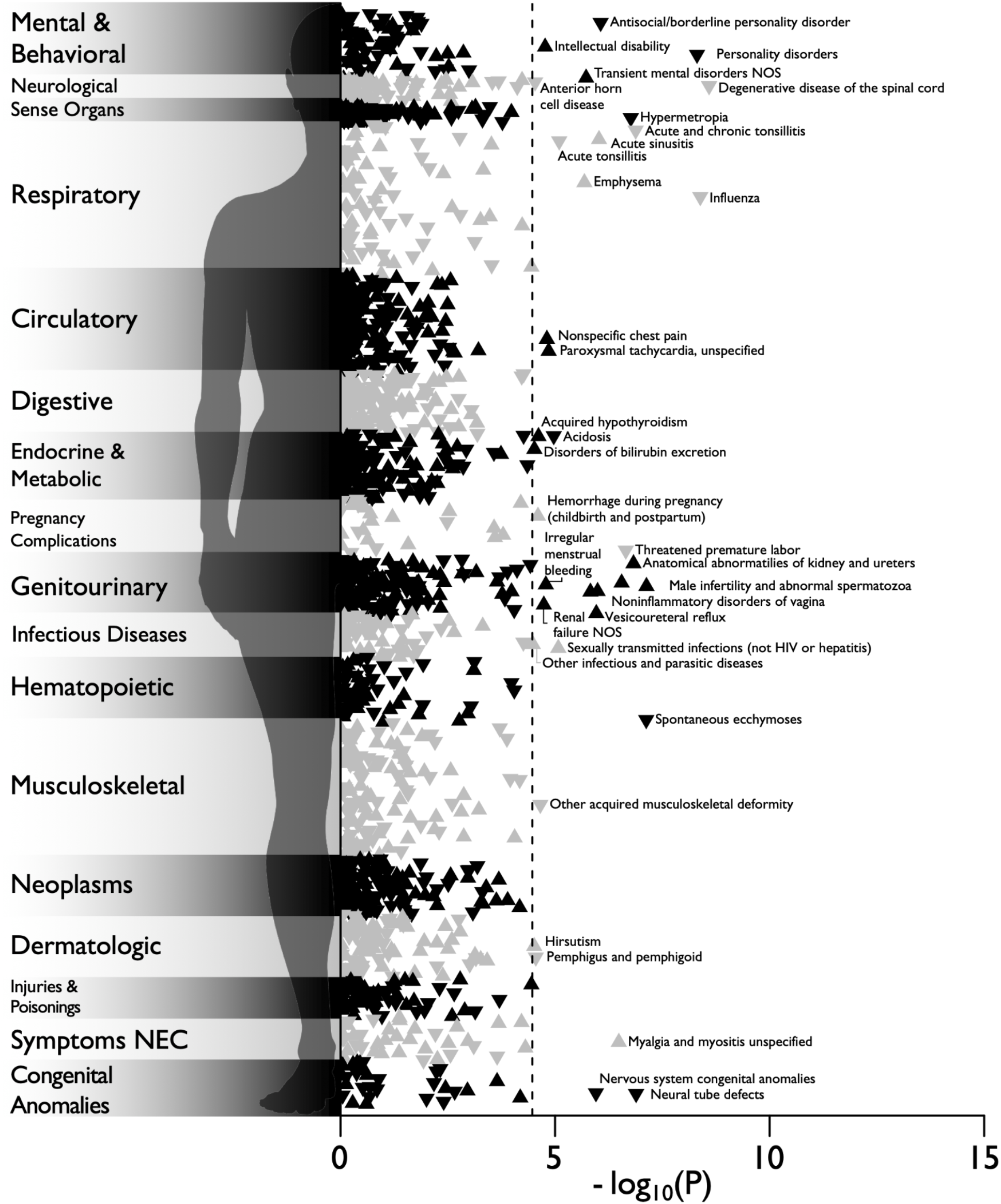
Phenome-wide association study (PheWAS) of a brain weight transcriptomic index in the Mass General Brigham (MGB) Biobank. Manhattan plot of the associations between brain weight transcriptomic index and 1,482 medical outcomes defined by “phecodes” in MGB Biobank, estimated with logistic regression in up to 37,272 patients. Associations are represented as triangles, where an upward-facing triangle denotes a positive relationship, while a downward-facing triangle denotes a negative relationship. The x-axis refers to the statistical significance of the association (plotted as -log_10_ P-values), the y-axis refers to the category of disease/disorder, and the dashed line denotes the P_Bonferroni_ < 0.05 significance threshold (i.e., 0.05/1,482 = 3.37e-5). Legibility necessitated that some condition labels be omitted from the plot. The complete results, including case-control record counts, effect sizes, and standard errors, are reported in Table S14.

## Discussion

Human brain size changes dynamically during the lifespan and varies substantially across individuals. Using available postmortem brain RNA sequencing and coupled ancillary data including brain weight (BW) from over two thousand human brain samples, we identified hundreds of genes that are differentially expressed across the BW continuum. Our results are replicable across independent datasets, show evidence of specificity to BW as compared to correlated traits such as height and weight, and reveal many BW genes that have not been implicated previously in imaging-genetic studies of brain size-related phenotypes. Overall, genes up-regulated in individuals with larger brains (BW+) or up-regulated in individuals with smaller brains (BW-) showed highly divergent spatio-temporal and cell-type-specific patterns of gene expression, and opposing differential expression patterns across species and in patients with neuropsychiatric disorders. These results pinpoint unreported molecular features of human brain size variation, and highlight the value of integrating measurements of brain morphology with postmortem transcriptomics.

Prior studies of brain size have primarily relied on *in vivo* neuroimaging, which have been only indirectly linked with brain gene expression because of the relative inaccessibility of in vivo human brain tissue. The burgeoning field of “imaging-transcriptomics” ^68–70^ has demonstrated a high degree of spatial alignment between neuroimaging “gradients” – representing topographical variation of brain morphology, function, or connectivity typically derived from magnetic resonance imaging (MRI) data ^71–73^ – and the spatial topography of gene expression identified in high-resolution transcriptomic resources such as the Allen Human Brain Atlas (AHBA) ^25,74–77^. However, only one previous study has directly examined brain size and brain gene expression using the comprehensive Allen Mouse Brain Atlas ^78^, finding a robust power-law scaling relationship between spatial gene expression gradients and brain size across mouse development – a model which accurately predicted human brain size ^79^. The present study extends these non-human results by demonstrating convergence between the spatial topographies of (1) BW-relative expression in the AHBA; (2) BW-gene set enrichment across neuroimaging GWAS; and (3) and nonlinear (hypo-allometric) scaling between regional and total surface area across individuals, primate species, across developmental stages ^26,27^. Collectively, these results suggest a direct link between neuroimaging gradients and the differential expression of BW genes.

The opposing developmental trajectories of BW gene sets, such that the expression of BW+ genes generally increases with age while that of BW-genes decreases with age, are highly suggestive of functional differences between these genes and their relationship with brain size. The difference in developmental trajectories of BW+ and BW-genes is corroborated by the anatomical convergence between the BW +/- relative expression map in AHBA and the topography of volumetric growth during human development ^4^. We find that brain areas with higher BW+ relative expression (frontal, parietal, temporal cortices) undergo protracted maturation with peak volume occurring later in life, while brain areas with higher BW-relative expression (insula, sensory cortices) achieve peak volume earlier in development. Moreover, the cross-species relative expression patterns in BW gene sets collectively mirror the expression differences observed in humans relative to non-human primates ^19^, with greater differences in the prenatal and late postnatal periods. BW-genes showed greater relative expression differences across both brain regions and cell types during the *prenatal* epoch – and these differences showed more pronounced expression in humans compared to macaques – which may be explained by BW-genes’ significant enrichment for molecular pathways involved in overall organismal growth and neurogenesis in particular. Notably, *ID4*, a BW-gene that functions as a transcriptional regulator, was previously implicated as part of a human-specific neural progenitor class showing earlier and higher expression in human fetal development compared to mice, and *Id4*-deficient mice also exhibit decreased brain size and mistimed neurogenesis ^80,81^. In contrast to BW-genes, BW+ genes showed relative expression differences across brain regions and cell types during *postnatal* periods (differences that were, again, more pronounced in humans relative to macaques). Of particular note, the BW+ cell membrane protein-encoding gene *FREM3* has been shown to have human-specific expression in deep cortical layer III glutamatergic neurons and distinct relationships with morphological and electrophysiological properties relative to other supragranular genes ^82^, pinpointing a potential role in the origination of long-range cortical feedforward projections that emerge during childhood ^83^.

The aforementioned spatiotemporal associations demonstrate the critical link between BW-genes and brain growth during dynamic periods of human neurodevelopment, well before peak brain size is reached. Thus, BW-genes can be interpreted as being both highly expressed *in* individuals with lower brain weight and *when* the brain is smallest. As only postnatal data were available to initially define the BW gene sets, the limited prevalence of high expression of BW-gene transcription in specific cell types later in life suggests that overall brain size is heavily influenced by critical periods of neuronal proliferation and migration when BW-genes are highly expressed. This hypothesis is in line with the finding of BW-relative expression differences in distinct postnatal cell types in humans relative to non-human primates, including inhibitory neurons in cortical layer IV. Notably, the AHBA BW +/- relative expression map and respective bio-annotations – against cytoarchitectonic, connectivity, and spatial expression data – further support the relationship between BW-genes and cortical layer IV. Previous work has demonstrated the potential role of protocadherins in the specification of layer IV identity in mice ^84^, and 8 protocadherin family genes were identified within the BW-gene set (and none in the BW+ set). This gene superfamily is involved with dendritic arborization and synaptic functioning, and has been implicated in a number of neuropsychiatric and neurodegenerative diseases ^85^.

There was direct evidence that BW genes are relevant to human disease. Intriguingly BW+ genes tended to be down-regulated, while BW-genes tended to be up-regulated, in individuals with schizophrenia and Alzheimer’s disease, two disorders that are robustly associated with decreased brain size ^4^. This trend was also seen in bipolar disorder and autism spectrum disorder, informing longstanding debates about the relevance of mechanisms that influence brain size to the pathophysiology of neuropsychiatric disorders in light of many shared genetic risk factors ^48,86,87^. Although these BW differential expression associations were identified in postmortem transcriptomic datasets, the translation of the cumulative BW-transcriptome score using imputed expression in the Mass General Brigham Biobank yielded associations to diverse clinical phenotypes that aligned with the bioinformatic annotations of BW gene sets. Notably, the relationship between the BW-transcriptome score and general mental health problems, intellectual disability, and neurodegenerative conditions helps validate the translational potential of this approach.

Of particular interest were phenotypic relationships with congenital brain-related anomalies, which were associated with a negative BW-transcriptome score. While a comprehensive examination of the relationship between gene dosage sensitivity and mechanisms that influence brain size is outside the scope of the current study, we did observe increased loss-of-function intolerance as well as enrichment for known causes of microcephaly in BW- (and not BW+) genes. Two of twelve genes (*PHC1, CEP135*) involved in primary microcephaly were within the BW-gene set ^88^, and recent work has also demonstrated the causal role of BW-genes *TRIM71* and *EGFR* in the pathogenesis of hydrocephalus-induced and viral-induced microcephaly, respectively ^89,90^.

The genetic regulatory findings from the TWAS/mmQTL analyses further highlight the epidermal growth factor (EGF) signaling pathway, which is critical for early brain development and has been implicated in myriad human diseases across the lifespan ^91–93^ – from psychiatric and neurodegenerative diseases to primary and non-primary brain cancers. For example, *EGFR* (a BW-gene) is one of the most commonly mutated genes found in glioblastoma ^94,95^, originating from neural stem cells in the subventricular zone ^96^, and recent single-cell sequencing work has identified *EGFR*-positive pre-oligodendrocyte progenitor cells as a novel intermediate progenitor cell type critical for human cortical expansion and myelination ^92^. Although postmortem transcriptomics necessarily prohibits within-subject longitudinal studies, there is a rich pharmacological history of *EGFR*-inhibitors ^97^ in the context of anti-cancer therapy and Alzheimer’s Disease ^98^, as well as proof-of-concept application of *EGFR* treatment in ameliorating brain growth in diffuse white matter injury in premature infants ^99^. Coupled with novel *in vitro* technologies (e.g., induced pluripotent stem cells, organoids), as well as the findings herein, this mounting body of work establishes an exciting opportunity for targeted experimental and interventional studies leveraging brain size as a critical outcome measure of development and aging.

Some methodological aspects of the present study deserve further consideration in future work. First, most donor samples came from the cerebral cortex (specifically dorsolateral prefrontal cortex). The GTEx dataset is an exception, with samples coming from numerous cortical and non-cortical regions; however GTEx contains considerably fewer donors than the PEC and ROSMAP datasets. Future work will incorporate additional brain regions to assess variability in differential gene expression with respect to variation in brain weight. Second, brain weight was used as a proxy for brain size in differential expression analyses. Although future studies could directly measure brain volume in addition to weight, the linear association between brain weight and volume has been clearly established. Finally, all donors for the differential expression analysis were sampled during the postnatal period. Given many of our results highlight the importance of prenatal epochs, future work using similar transcriptomic data in fetal tissue will be critical for comparative analysis during early periods of neurodevelopment.

Despite these methodological considerations, the present study comprehensively outlines the transcriptomic underpinnings of human brain size variation and demonstrates the importance of postmortem measurement of brain size as a quantitative trait to incorporate into transcriptomic analyses. Examined separately or in contrast to one another, the BW+/- gene sets define the spatiotemporal extremes of brain gene expression, charting a molecular segregation of brain regions and developmental epochs. Our results implicate BW genes in the expansion of the human brain relative to non-human primates, as well as gene expression differences observed in brain tissue from patients with neuropsychiatric and neurodegenerative disorders with established differences in brain size. The BW-transcriptome score bridges *ex vivo* transcriptomics and *in vivo* genetic risk, highlighting the role of BW genes in both primary and non-primary brain conditions. With ever-increasing data from open science initiatives, parallel innovations in RNA sequencing technology, and widespread utility of clinical genetics, the power of leveraging phenotypic associations from readily accessible transcriptomic datasets holds tremendous promise for future translational studies of myriad human traits and diseases.

## Acknowledgements

The authors would like to acknowledge the various open science initiatives, and their associated funding, that provided the necessary datasets used in this manuscript. JS and JWV were supported by NIMH T32MH019112. AFA-B and JS were supported by NIMH K08MH120564. TTM was supported by NHGRI T32HG010464. VW was supported by St. Catharine’s College Cambridge. RAIB was supported by the Autism Research Trust. KW was supported by the Wellcome Trust (215901/Z/19/Z). PEV is a Fellow of MQ: Transforming Mental Health (MQF17_24). TDS was supported by R01MH113550, R01MH120482, R01MH112847, R01EB022573, RF1MH121867, and R37MH125829. GB was supported by the Australian National Health and Medical Research Council (NHMRC; Investigator Grant 1194497). We thank Mass General Brigham Biobank for providing genomic and health information data.

## Author Contributions

Conceptualization: J.S., A.F.A.-B. Methodology: J.S., T.T.M., M.J.G., A.F.A.-B. Formal Analysis: J.S., T.T.M., Y.H.L., L.M.H., S.T., B.N., C.G., T.G., M.J.G. Resources: J.S., T.T.M., V.W., R.A.I.B, J.W.S., T.G., M.J.G., A.F.A.-B. Data Curation: J.S., T.T.M., J.W.V., Y.H.L., V.W., R.A.I.B., S.T., B.N., C.G., T.G., M.J.G. Writing – Original Draft: J.S., T.T.M. M.J.G., A.F.A.-B. Writing – Review & Editing: all authors. Visualization: J.S., T.T.M. Supervision: M.J.G., A.F.A.-B. Project Administration: J.S., A.F.A.-B. Funding Acquisition: J.S., A.F.A.-B.

## Methods

All analyses in this manuscript make use of available datasets and resources, which are summarized in table below. Data and code for analyses will be made available online (https://github.com/jms290/brain-weight) upon publication.

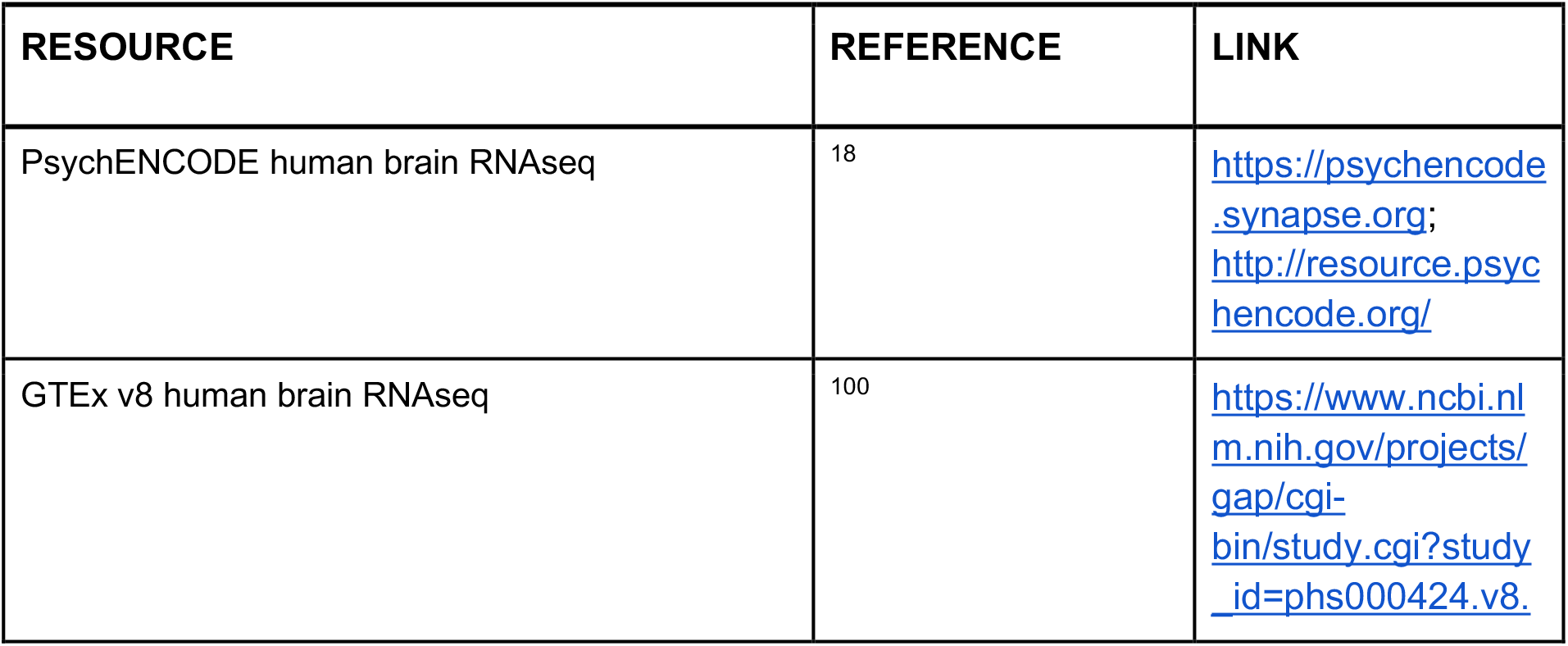
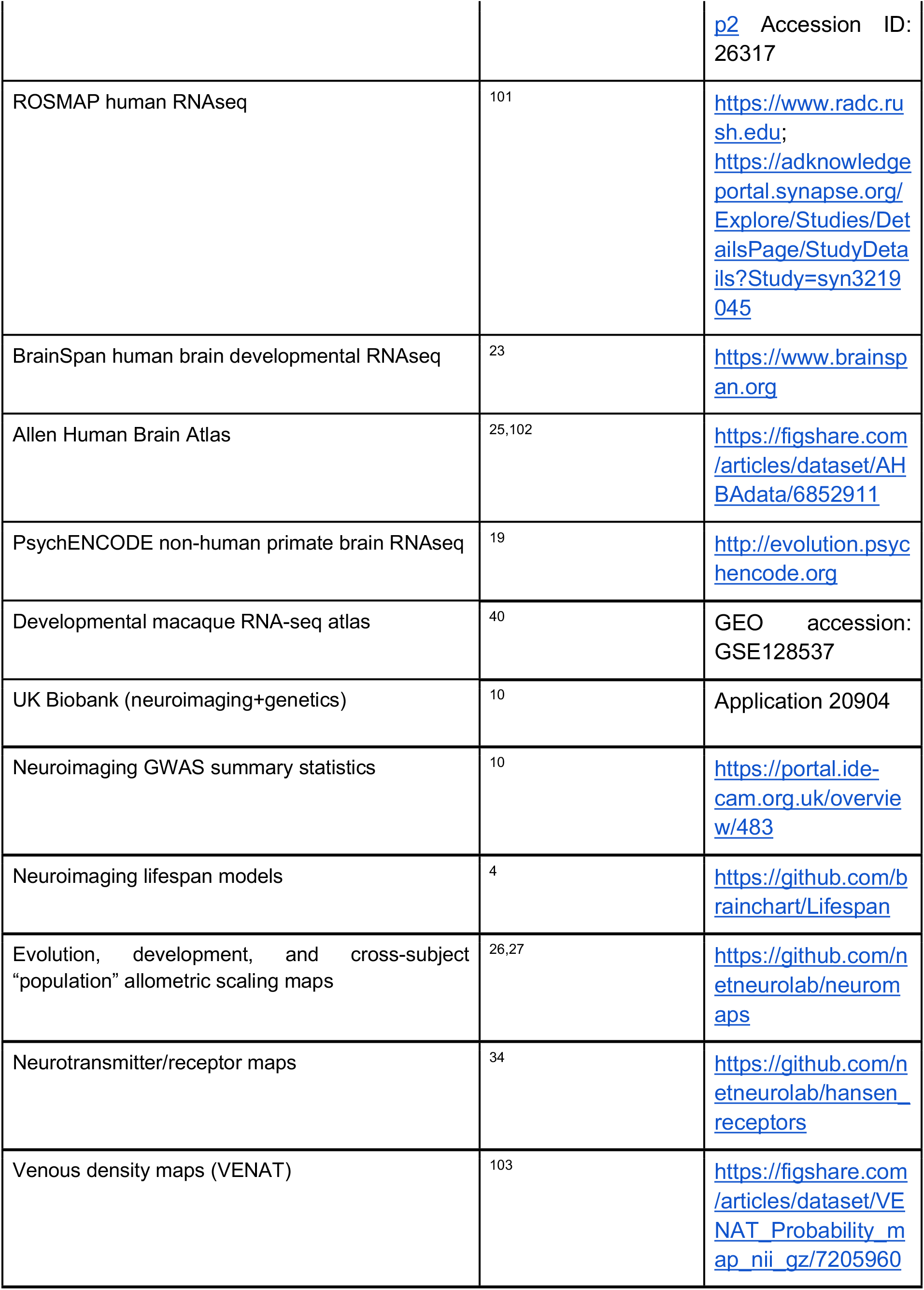
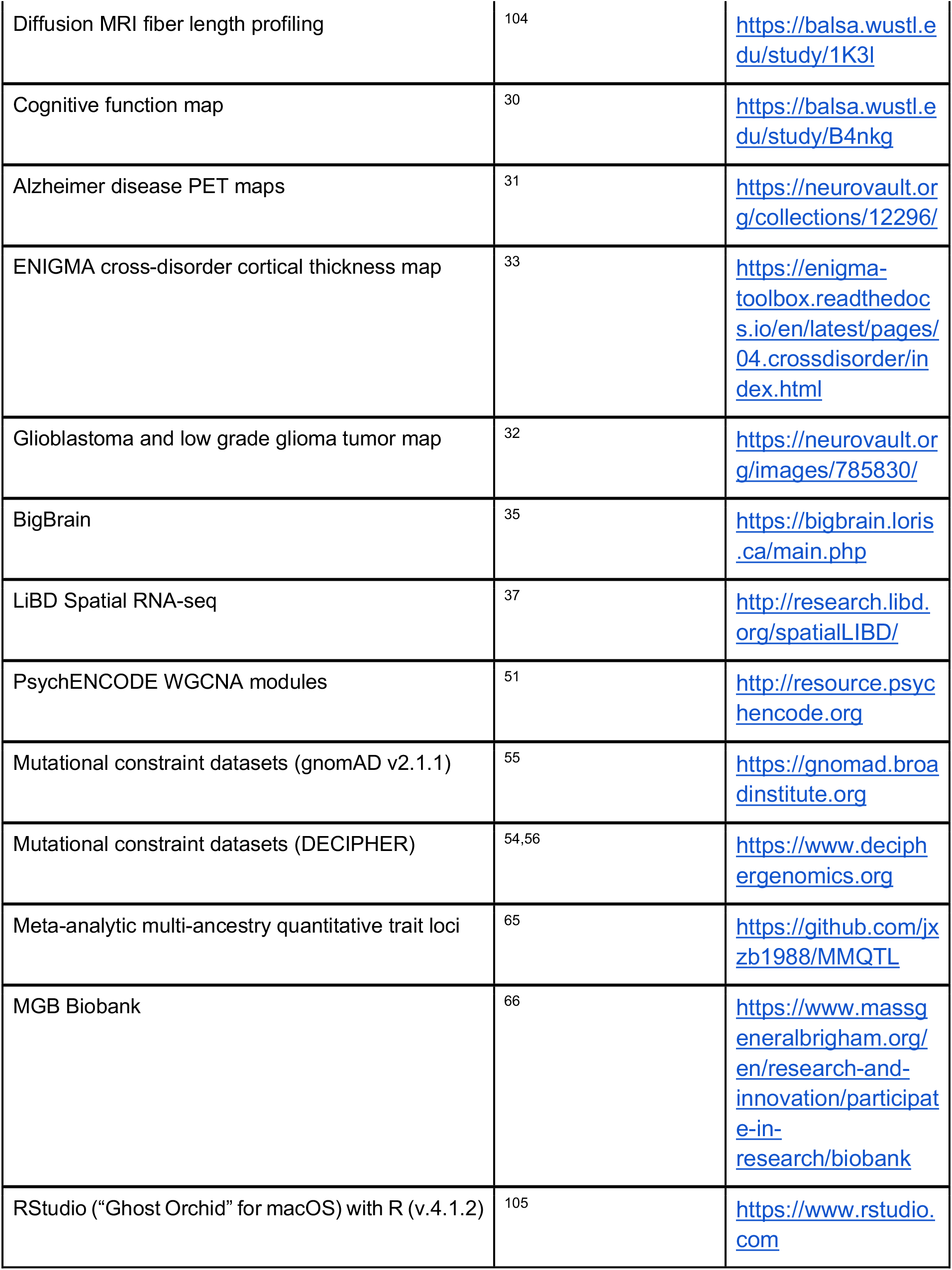
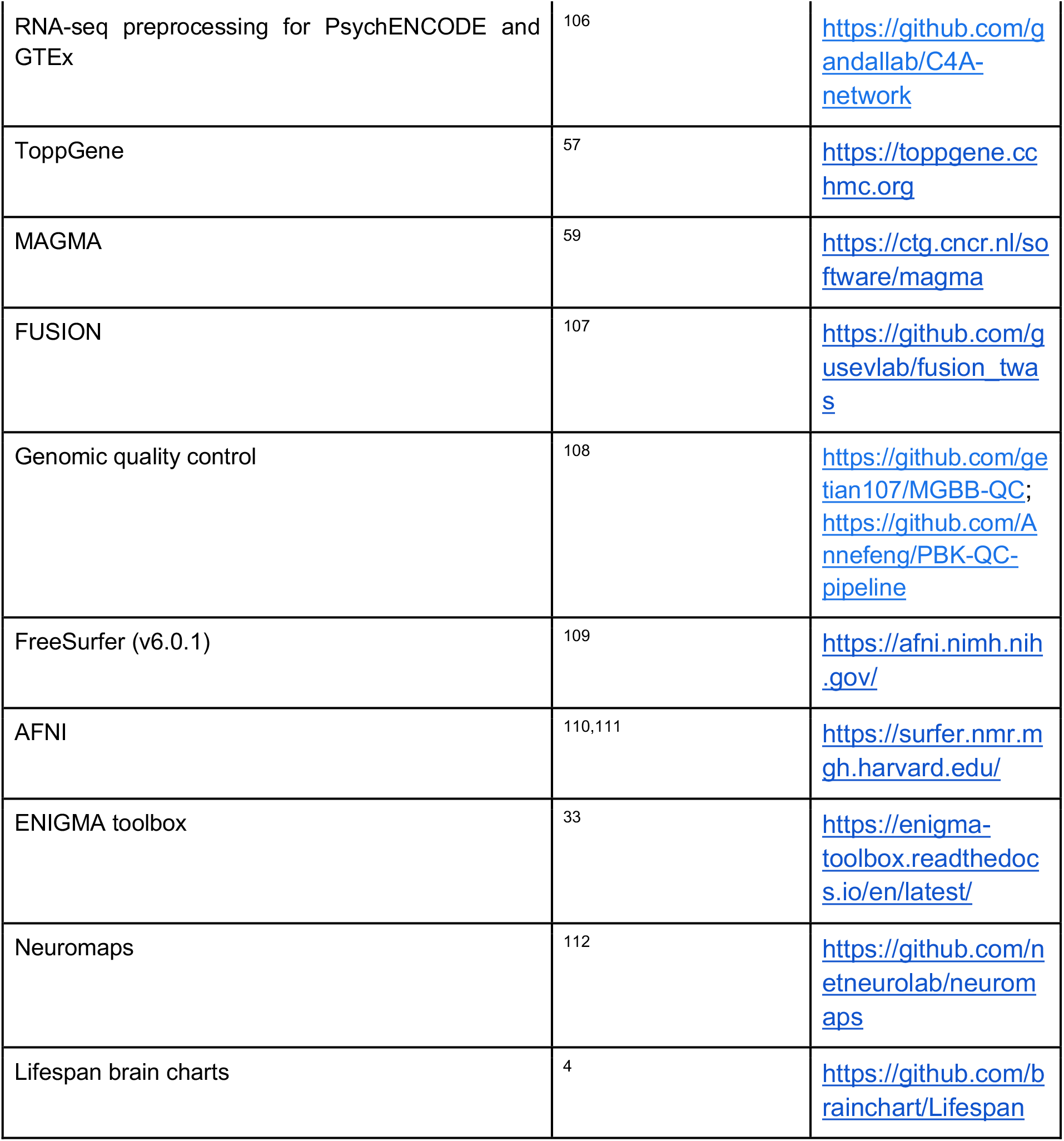

### Validation of brain weight predictions from lifespan neuroimaging models

Brain weight data (total N=3,689) from the three primary transcriptomic datasets (N=2,531; PsychENCODE, GTEx, ROSMAP) was combined with additional brain weight data from the literature ^113^. Metrics of brain size (total cerebrum volume and total cortical surface area) for each subject were predicted using life-spanning models of the two neuroimaging phenotypes ^4^. Subsequently, brain weights for each subject were then converted to volumes based on previous estimates of average brain density ^20^, and correlations were computed between converted brain weight and predicted brain size (volume or area).

### Brain weight differential gene expression analysis

All transcriptomic datasets were downloaded from accessible repositories listed in the table above, with methods described previously in the respective source citations. Briefly, the PsychENCODE “freeze 2” dataset consisted of uniformly processed data from six studies: BipSeq, LIBD_szControl, CMC_HBCC, CommonMind, BrainGVEX and UCLA-ASD ^106,114^. Post-quality-control RNA-seq reads were previously aligned to the hg19 reference genome with STAR 2.4.2a and gene-level quantifications were calculated using RSEM v1.2.29. Genes were filtered to include those with >0.1 TPM in at least 25% of samples. Similarly, GTEx RNA-seq reads were aligned to the hg19 reference genome with STAR 2.4.2a and transcript-level counts quantified with RSEM v1.2.22. Samples from non-brain tissues and tissues with different sample preparation (cortex and cerebellar hemisphere) were removed. Additionally, samples with a history of disease possibly affecting the brain prior to filtering for features with CPM > 0.1 in at least 25% of samples were also removed. Gene-level counts were then normalized using TMM normalization in edgeR and log2-transformed to match PsychENCODE. Each brain region was then assessed for outlier samples, defined as those with standardized sample network connectivity Z scores < −3, which were removed. For the ROSMAP cohort, normalized data from previous publications was downloaded from the link provided in the table above based on previous work ^101^. RNA-seq data was aligned by Tophat v2.0 and v2.1 and transcript enrichments were estimated with RSEM. Quality metrics were provided by Picard, which was also used to mark duplicate reads. Within-batch normalization was conducted through quantile normalization while the between-batches normalization was conducted through ComBat ^115^. Overall, 25,774 genes were included in the PsychENCODE analyses after filtering, 21,347 in GTEx (19,481 overlapping with PsychENCODE), and 15,083 in ROSMAP (all overlapping with PsychENCODE).

For the PsychENCODE and GTEx cohorts, code for performing the transcriptomic normalization and differential expression analysis followed a previously published approach ^51,106^ with the addition of brain weight as an independent predictor variable (https://github.com/gandallab/C4A-network). Per gene linear models were implemented for PsychENCODE, and linear mixed effects models using the ‘lme4’ package in R were implemented for the GTEx cohort with the addition of a random effect for donor (to account for the multiple brain regions per donor). P-values for the linear mixed effects models were calculated using the likelihood ratio test, as implemented in the ‘lmerTest’ package in R. For ROSMAP, as for the GTEx dataset, per gene linear mixed effects models were implemented, including covariates used for the normalization, brain weight, and a random effect of study. Cross-cohort meta-analyses were performed at the level of gene-brain weight association statistics (beta coefficients) and empirical p-values using the ‘metap’ packing in R.

### Neuroimaging data

Structural minimally processed (https://biobank.ctsu.ox.ac.uk/crystal/crystal/docs/brain_mri.pdf) T1 and T2-FLAIR weighted data was downloaded from the UK BioBank, and further preprocessed with Freesurfer (v6.0.1) ^109^ using the T2-FLAIR weighted image to improve pial surface reconstruction. The ‘recon-all’ reconstruction included bias field correction, registration to stereotaxic space, intensity normalization, skull-stripping, and white matter segmentation. When no T2-FLAIR image was available, Freesurfer reconstruction was done using the T1 weighted image only. Given systematic variation related to the inclusion of T2-FLAIR, this was included as a confound variable in downstream analyses. Following reconstruction, the Human Connectome Project multimodal parcellation “HCP-MMP” ^116^ was aligned to each individual freesurfer average image and parcellated values were extracted. Reconstruction reliability was assessed using the Euler index ^117^ and included as a covariate in subsequent analyses.

Structural diffusion weighted imaging was obtained for both datasets. In addition to MD and FA, we ran Accelerated Microstructure Imaging via Convex Optimization (AMICO) to estimate neurite orientation density and dispersion indices ^118^. The T1 aligned parcellation template was co-registered to the diffusion weighted image using fsl FLIRT and regional values for FA, MD and the three NODDI parameters were extracted using AFNI’s 3dROIstats function. Total sample size of the imaging dataset (to match with imputed expression data for the transcriptome-wide association analyses, described below) was 22,387 subjects.

### Neuroimaging transcriptome-wide association studies

To identify genes whose cis-regulated expression is associated with global and regional neuroimaging phenotypes in the UK Biobank *in vivo* dataset, we performed a series of transcriptome wide association studies (TWAS) leveraging individual-level genotype and postmortem brain expression data from PsychENCODE, described preiovusly ^51,107^. Briefly, TWAS was implemented using the FUSION software package (https://github.com/gusevlab/fusion_twas; ^107^) with custom SNP-brain-expression weights generated using the PsychENCODE dataset of 1321 unique individuals with imputed genotypes. Using the AI-REML algorithm ^119^ implemented in GCTA ^120^ implemented by the FUSION package, we first identified the subset (N = 14,750) of total expressed genes found to have significant cis SNP-heritability in our dataset (cis-h^2^_g_ uncorrected P < 0.05 within 1 Mb window around the gene body). SNP-expression weights were calculated in a 1Mb region around all heritable genes using expression measurements adjusted for diagnosis, study, age, age^2^, RIN, RIN2, sex, tissue, PMI, 20 ancestry PCs, and 100 hidden covariates ^51^. Accuracy of five expression prediction models were tested (best cis-eQTL, best linear unbiased predictor, Bayesian linear mixed model, Elastic-net regression, LASSO regression) using the most accurate model for final weight calculations as implemented in the FUSION package. TWAS neuroimaging-association statistics were computed using these custom weights, LD structure calculated from our PsychENCODE samples’ genotypes, and neuroimaging data from UK Biobank as described above. For each global and regional neuroimaging phenotype, TWAS association statistics were Bonferroni-corrected for multiple comparisons (final count N = 13,421 genes). At loci (+/- 100 kb) with multiple significant associations, joint and conditional association analyses were further performed as implemented in the FUSION.post_process.R script. Gene weights are available from http://resource.psychencode.org/.

### Bioinformatics analyses

#### Spatiotemporal annotation

Spatially-comprehensive cortical microarray gene expression data from the Allen Human Brain Atlas (AHBA) was processed and mapped to the same HCP-MMP atlas (N=180 regions) as the previously described neuroimaging data ^25,102^ for the purpose of assessing regional expression and relative expression differences in brain weight gene sets. Three regions (“MT”, “a10p”, “RI”) were removed due to inadequate sampling across donors and quality control. AHBA data was also mapped to the Desikan-Killiany atlas (N=34 regions) depending on the comparative maps used for spatial correlation. Additional multimodal neuroimaging, cytoarchitectonic, and functional maps (each projected to the same abovementioned cortical parcellations) can be found in the Methods. Developmental RNA-seq data as well as prenatal laser microdissection microarray data from BrainSpan ^23^ was used for mapping age trajectories and assessing early regional relative BW expression differences (see also Methods).

#### Cross-species effects

To assess relative expression differences in BW across species, we leveraged a matched human and non-human primate dataset associated with the PsychENCODE project ^19^. This dataset combines tissue samples from the six adult humans, five adult chimpanzee brains and five adult macaque brains. The dataset can be downloaded from http://evolution.psychencode.org where it is labeled as “Adult human, chimpanzee, macaque data” in the “Processed Data”/”mRNA-seq” tab. The dataset is pre-harmonized to include a consistent set of 16 cortical and subcortical brain regions with mRNA sequencing performed on 11,346 curated homologous genes. Similarly, to examine species-by-development effects, we used a second pre-curated dataset, downloadable from the same link, labeled “Developmental rhesus and human data”. This dataset includes tissue samples from 36 human brains (15 female, ages 8 post-conception weeks to 40 years, mean = 97 +/- 147 post-conception months) and 26 macaque brains (8 female, ages 60 post-conception days to 11 years, mean = 36 +/- 46 post-conception months) across 16 homologous cortical, subcortical and cerebellar brain regions. Three transient developmental brain regions were excluded (lateral, medial and caudal ganglionic eminence), while other prenatal regions were considered equivalent to their most similar adult brain regions (e.g., dorsal thalamus to mediodorsal thalamus, upper rhombic lip to cerebellum, etc). This was only relevant for two of 62 total brains that possessed these early developmental regions. Each brain region had mRNA sequencing for 27,932 genes.

Additionally, to assess the consistency of regional BW +/- relative expression differences across species, we used an independent postmortem brain gene expression dataset (RNAseq; see Methods) in macaques containing 416 samples from 8 macaques across 52 cortical and non-cortical brain regions ^40^. Processed data was available (see Methods), and we included biological (age, sex) and technical (RIN, medTIN) covariates when examining BW +/- relative expression differences for each brain region. Each brain region had mRNA sequencing for 14,898 genes.

#### Cell-type relative expression differences

To assess relative expression differences in BW across cell types, we leveraged aligned developmental single cell (fetal) and single nucleus (adult) RNA-seq datasets in both humans and non-human primates. Macaque data was downloaded from http://evolution.psychencode.org, where they are labeled as “Fetal expression matrix” and “Adult expression (Rdata)” under the “Processed Data”/”Single cell RNA-seq” or “Processed Data”/”Single nucleus RNA-seq” tabs, respectively. These datasets contain expression data on ∼15k genes across various cell types sampled in the dorsolateral prefrontal cortex (DFC) from two (fetal) and three (adult) macaques. More detailed protocol information can be found in previous work ^19^.

Human data were downloaded from http://development.psychencode.org, where they are labeled as “scRNA-seq (Rdata)” and “snRNA-seq (Rdata)” under the “Processed Data”/”Single cell/nucleus RNA-seq” tab. For over 20k genes, the fetal single cell dataset contains expression data sampled across the pallium (primarily labeled as “DFC”) from nine donors, and the adult single nucleus dataset contains expression data in the DFC from three donors. More detailed protocol information can be found in previous work ^18^.

#### Gene ontology enrichment

Brain weight gene lists were submitted as gene sets to ToppGene’s ToppFun enrichment feature (https://toppgene.cchmc.org/enrichment.jsp). The following term categories were assessed: GO: Molecular Function, GO: Biological Process, GO: Cellular Component, Pathway (all), Mouse Phenotype and Human Phenotype. All other settings were left to their defaults. Note that ToppGene databases are continuously updated; this ToppGene query was last conducted on October 23rd, 2021.

#### Functional enrichment

The gene set enrichment analysis further nominated associations with brain morphological phenotypes derived from in vivo structural magnetic resonance imaging (MRI). To validate this finding, we used an established gene-set analysis method, MAGMA ^59^, to evaluate the enrichment of BW gene sets against genome-wide association studies of global and regional neuroimaging phenotypes ^10^, as well as metrics of head size – head circumference and intracranial volume ^62–64^.

### Phenome-wide association study in Mass General Brigham Biobank

Phenome-wide association study (PheWAS) analyses were performed in the Mass General Brigham (MGB) Biobank, a biorepository from the MGB healthcare system based in the greater Boston area with patient data on electronic health record, genetic, and lifestyle variables ^66^. Launched in 2010, the MGB Biobank has enrolled 138,042 participants and generated genotyping microarray data for more than 65,265 participants to date. To reduce the risk of population stratification, we restricted all PheWAS analyses to 37,272 patients of European ancestry (22,232 were genotyped using Illumina MEG, MEGA, or MEGA EX arrays and the remaining 15,040 were genotyped using the Illumina GSA array). The recruitment strategy, genotyping procedures, and quality control procedures of MGB Biobank have been described previously (see Methods) ^66^. All participants provided written consent upon enrollment. MGB Institutional Review Board approval for the present analyses is covered under protocols #2009P002312 and #2021P003641.

We used the FUSION package in R, as in the TWAS, to impute individual-level gene expression for patients in MGB Biobank, using pre-computed weights from the PsychENCODE consortium as the reference transcriptome dataset (gene weights available from http://resource.psychencode.org/). We then calculated a cumulative brain weight transcriptomic index as a coefficient-weighted sum of imputed gene expression – similar to how polygenic indexes are calculated using single nucleotide polymorphism data. This score was subsequently standardized and linked to electronic health record data from the MGB Biobank participants. Case status for each medical phenotype was assigned using a standard “phecode” approach ^121^, where the presence of at least two International Classification of Disease (ICD)-10CM codes was required. PheWAS was then conducted using the PheWAS package in R (https://github.com/PheWAS/PheWAS), fitting logistic regression models to each of the 1,482 medical outcomes in order to estimate the odds of each diagnosis given the brain weight transcriptomic index while accounting for sex assigned at birth, current age, genotyping batch, and the first 10 genetic principal components as covariates.

### Statistical analyses

For all analyses in this study significance was determined at P < 0.05, after Bonferroni correction for multiple comparisons (noted P_Bonferonni_). For the brain weight discovery analyses in PsychENCODE this refers to an uncorrected P < 1.94e^-6^ threshold. For the TWAS analyses, P_Bonferroni_ < 0.05 equated to an uncorrected P < 3.73e^-6^ for global and uncorrected regional (surface-based: P < 2.07e^-8^, volume-based: P < 1.19e^-8^) phenotypes. False Discovery Rate (FDR) correction was used in lieu of Bonferroni correction to determine significant brain weight gene sets (P_FDR_ < 0.05) in the intersection analyses with the differential expression results across disorders based on how these data were thresholded in their original publications. Additionally, where applicable, we used two different permutation-based tests for empirical statistics to complement uncorrected P-values based on context-specific robust null models (P_permutation_ and P_spin_), as implemented previously ^122^. P_permutation_ denotes performing gene-level statistical analyses using 10,000 randomly sampled gene sets (without replacement) of similar size to the empirical sets, drawn from the entire gene list used for the brain weight differential expression analysis in PsychENCODE. P_spin_ denotes performing spatial correlations between two brain maps using 10,000 “spins” (preserving spatial autocorrelation) of one map (see also Methods). The term “relative expression” implies the use of Student T-tests comparing BW+ vs. BW-gene sets. Unless otherwise noted, all P-values for linear mixed models were calculated using likelihood ratio tests as implemented in the “lmerTest” package in R. As this was a retrospective study, no statistical methods were used to pre-determine sample sizes, however, collectively, this study makes use of the three largest datasets of postmortem human brain tissue. As such, we comprehensively assessed the main brain weight differential expression results explicitly in a discovery (PsychENCODE) and in two independent replication datasets (GTEx and ROSMAP). Randomization and blinding were not possible due to the study being retrospective and observational. Accordingly, subject-level covariates were used to account for variation in gene expression as well as to remove unwanted confounding effects. Normalized gene expression was assumed to follow normal distribution, but this was not formally tested. Additional details for each analysis are provided in relevant sections above.

## Data and code availability

All data and code for this paper can be found in the information provided in the Methods.

## Notes

### Competing Interest Statement

The authors have declared no competing interest.

