## Supplemental Material for "The molecular genetic landscape of human brain size variation"

### Supplemental information

#### Replicability of the brain weight transcriptomic relationships

We performed additional assessments of replicability In addition to the meta-analytic approach for replicating the gene models from PsychENCODE in two independent datasets, Genotype-Tissue Expression (GTEx) and Religious Order Study/Memory and Aging Project (ROSMAP). Rank-rank hypergeometric overlap (RRHO) ^1^ analysis confirmed significant replication in both datasets: GTEx (N = 227 subjects; Spearman r = 0.39, P < 0.0001; 417/846 significant overlapping genes) and ROSMAP (N = 634 subjects; Spearman r = 0.34, P < 0.0001; 525/847 significant overlapping genes).

#### Influence of age on the relationship of brain weight and gene expression

Considering the clear difference in expression trajectories, we performed a post hoc differential expression analysis across genes to examine the interaction between BW and age. 222 genes showed significant (P_Bonferroni_ < 0.05) BW-by-age interaction effects (linear or quadratic age terms) without showing either significant age or BW effects (both P_Bonferroni_ > 0.05) – 130 genes with positive coefficients denoting the effect of BW on expression increasing with age and 92 genes with negative coefficients denoting the opposite pattern (**Fig. S4B**).

#### Transcriptome-wide association studies of neuroimaging phenotypes in the UK Biobank

##### Heritability, genetic and phenotypic correlations

Consistent with prior reports, heritability of phenotypes derived from our image processing pipeline was moderate-to-high, supporting downstream tests of association with imputed expression of specific genes. Total SNP-based heritability (*h*^2^_SNP_) for global neuroimaging phenotypes was calculated by genome-wide complex trait analysis (GCTA) ^2^, using common typed and imputed variant (MAF > 0.01, genotyping call rate > 0.95, Hardy-Weinberg p-value > 1x10^-6^) SNP data from 20,387 individuals. We found the highest *h*^2^_SNP_ for total surface area (SA; h^2^ = 0.57, SE = 0.03, P< 1e^-12^) and the lowest with mean fractional anisotropy (FA; h^2^ = 0.16, SE = 0.03, P = 1.88e^-10^), with the size (e.g, SA) and shape (e.g. curvature) metrics showing higher heritability values than the microstructural diffusion imaging features related to composition (**Fig. S16**).

There was a clear correlation structure between imaging phenotypes in terms of their pattern of association with imputed gene expression (rho_GE), which notably diverged from the phenotypic correlation between imaging phenotypes (rho_p) in some cases. For each pair of imaging phenotypes, rho_GE was calculated as Spearman’s correlation of the transcriptome-wide association study (TWAS) P-values across all genes. Some inter-phenotypic relationships were consistent between rho_p and rho_GE as predicted from prior work: high positive correlation between volume and SA, negative correlation between CT and SA, and positive correlation between different indices of cortical curvature. However, strong rho_GE associations also emerged which were not present in rho_p, for instance high rho_GE between mean diffusivity (MD) and neurite orientation dispersion (OD) and a greater preponderance of negative correlations in rho_GE compared to rho_p overall (**Fig. S16**). This suggests that TWAS pleiotropy, wherein imputed expression of a single gene is associated with variance in multiple MRI metrics, does not simply recapitulate patterns of inter-individual covariance in the underlying phenotypes.

##### Global and regional neuroimaging TWAS

At the global level, TWAS identified 39 associations (29 genes, 25 unique LD clusters) wherein genetically imputed brain expression was associated with at least one of the twelve neuroimaging phenotypes at genome-wide levels of significance (P_Bonferroni_ < 0.05; **Table S5**). There were 8 genes significantly associated with average folding or thickness, 3 genes associated with average diffusion metrics, and 13 genes associated with total volume or surface area. In addition, there were 5 genes that influenced multiple phenotypes across these categories (*EGFR,* *EML2, PCCB, SUOX, TFDP2*). The percent variance explained, quantified as the partial correlation after controlling for other covariates, ranged from 3% to 6%. We found no significant interaction effects of sex or age on the imputed expression of any gene for any of the neuroimaging phenotypes (all P_Bonferroni_ > 0.05). Fine-mapping was used to identify high probability genes within LD blocks (**Table S12**).

At the regional level when controlling statistically for the linear relationship with the global phenotypes, there were 265 significant associations (27 folding/thickness, 106 diffusion, 132 volume/surface area; **Table S7**). Regional associations implicated a total of 123 unique genes and 92 unique LD clusters. These regional associations were robust to sensitivity analyses i) controlling for a non-linear relationship between global and local phenotypes and ii) not including the global phenotype in statistical models (**Table S8**). For the significant regional associations, the percent variance in imaging phenotypes explained by imputed gene expression ranged from 4% to 11%, and there were no significant interaction effects for sex or age. All significant gene-phenotype were tested for significance at uncorrected P < 0.05 threshold using a discovery/replication breakdown of 90/10 percent, respectively. Cumulatively, 119 associations replicated in contrast to 16 which would be expected to replicate by chance.

Cumulatively, our results include five genes – *EGFR*, *LAMC3*, *GINM1*, *STH* and *LRP11* – with significant associations with both global and regional imaging phenotypes. We investigated the possibility that our conservative threshold for genome-wide significance could mask evidence of additional global-regional pleiotropy. For the significant regional associations, we therefore performed a post-hoc assessment of the level of support for an association between the same genes and the corresponding global phenotype. Out of the 265 significant regional associations, there was a trend towards a global association (uncorrected P < 0.01) in only 60 cases, supporting the theory that the large majority of genes with local genetic influences may not influence global imaging phenotypes.

In addition, we investigated whether there was concordance in the direction of regional effects compared to global effects, i.e. whether the statistical effect was in the same direction for gene-phenotype pairs. Overall, there was a moderate tendency towards more concordance, with 182 out 265 (69%) of effects being in the same direction. This tendency was accentuated when only considering gene-phenotype pairs for which there was at least trend-level support of a global relationship as described above, for which 48/60 (80%) of effects were concordant. These results support the theory that regional effects represent a mixture of i) localized attenuation of a global effect, ii) localized augmentation of a global effect, and iii) truly local effects without a global corollary.

Finally, we investigated whether there was evidence of discordance in the direction of regional effects themselves. Given the high degree of covariance between brain regions, we expected most if not all regional effects to be concordant in terms of the direction of effects. Strikingly, although most regional associations were concordant for a given gene, there was strong evidence of discordance for 8 genes in regional effects that reached genome-significance. Two of these genes (*CEP120* and *KRT8P33*) affected regional diffusion metrics, although as these genes are adjacent on chromosome 5 the true effect is likely limited to a single genomic locus. Additionally, two genes (*ZIC4* and *DEPDC1B*) affected regional estimates of surface area, while four genes (*NDUFAF2*, *METTL10*, *PARPBP* and *EFCAB5*) affected regional estimates of volume. As this discordance persisted whether or not global phenotypes were included as covariates, the discordance does not appear to merely reflect relative deviations from global effects. Indeed for the 6 genes affecting surface area or volume, there is little evidence of a consistent global effect (all P > 0.01).

Regional discordance in local genetic effects could be constrained by known patterns of regionally-specific brain development. To investigate this possibility, we examined the allometric scaling coefficients of discordant brain regions. Greater hyper-allometry (scaling coefficient > 1) is indicative of increased local cortical expansion relative to total brain size, while greater hypo-allometry (scaling coefficient < 1) is indicative of decreased local cortical expansion relative to total brain size. Moreover, there are known evolutionary underpinnings to regional variation in allometric scaling of surface area and volume. Interestingly, we observed that discordance in regional genetic effects appeared to be associated with regional differences in allometric scaling (average difference in scaling coefficient between discordant regions = 0.30; permutation test P_spin_ = 0.0053). This association – between allometric scaling and locally distinct genetic effects – implicates a set of genes with divergent effects on brain areas that have undergone relatively recent expansion in human evolution compared to non-human primates.

##### Overlap with prior neuroimaging-genetics studies

The large majority of genes identified in the present study have previously been nominated gene-mapping of significant SNPs from prior neuroimaging GWAS studies, based on queries of GWAS catalog (<https://www.ebi.ac.uk/gwas/>) and the latest UKBB GWAS (<https://open.win.ox.ac.uk/ukbiobank/big40/>). The present results provide more direct evidence of an association between imaging phenotypes and the genetic regulation of expression patterns of these genes in the human brain. Of note, only 15% of genes we report overlap with neuorimaging-associated genes from a recent cross-tissue TWAS ^3^. Although more than expected due to chance (hypergeometric test, P = 0.005), this relatively small overlap highlights the importance of methodological differences in the approach to imputation in TWAS analysis (see Discussion). A subset of 12 significant genes in our discovery analysis have not previously been reported to be associated with neuroimaging phenotypes to our knowledge, however, none of these genes were statistically significant in the replication analysis (see Supplement).

Of potential interest is the subset of genome-wide significant genes that have not previously been associated with imaging phenotypes, although as noted in the main text these associations were not significant in the replication analysis. Based on queries of GWAS catalog (<https://www.ebi.ac.uk/gwas/>) and the latest UKBB GWAS (<https://open.win.ox.ac.uk/ukbiobank/big40/>), the following genes have not previously been reported to be associated with neuroimaging phenotypes: *ATG14, AXIN2, GCDH, HAGH, IARS2, INPP5F, KCNG1, KCTD9P1, KIAA1161, MYRIP, NMB, RNF170*, *SERPIND1*. This subset of genes included associations with global imaging phenotypes (*HAGH, IARS2, KIAA1161, MYRIP, INPP5F*) and regional imaging phenotypes (*ATG14, AXIN2, GCDH, KCNG1, KCTD9P1, NMB, RNF170, SERPIND1*). They also included associations with size (*INPP5F, KCNG1, GCDH, ATG14, AXIN2, KCTD9P1*), shape (*KIAA1161, MYRIP, IARS2, HAGH*) and composition (*SERPIND1, RNF170, NMB*). Some of these genes have previously been mapped to SNPs associated with GWAS neuropsychiatric disease. For example, *NMB*, which in the present study is most strongly associated with neurite orientation dispersion in primary motor cortex, has previously been implicated in GWAS of schizophrenia and bipolar disorder ^4^; and *INPP5F*, which is most strongly associated with total cortical surface areas in the present study, has been implicated in GWAS of Parkinson’s disease ^5^. Of note, one of these genes (*SERPIND1*) was also reported in a recent cross tissue neuroimaging TWAS in UKB ^3^.

##### Overlap with brain weight transcriptomic results

To examine the genetic regulatory aspect of the brain weight (BW) transcriptomic findings, we assessed the degree to which BW gene sets overlapped with significant genes across neuroimaging TWAS. In general, BW genes significantly overlapped with 29 transcriptome-wide significant global (P_permutation_ < 0.01) and 124 unique transcriptome-wide regional (P_permutation_ < 0.05) genes associated with multimodal neuroimaging phenotypes (using a TWAS-study threshold of P_Bonferroni_ < 0.05 across regions and phenotypes to establish significant gene sets). Specificity analysis across neuroimaging phenotypes did not reveal any significant differences in BW gene set overlap or enrichment.

#### A polygenic transcriptome score for brain weight shows disease-related differences

Based on the overlap of BW gene sets with differentially expressed genes in patients with neuropsychiatric disease, we developed a BW transcriptome score – calculated for each individual as the linear combination of weighted gene expression (i.e., sum of expression across genes weighted by the PsychENCODE BW model coefficients). Using all available genes, this cumulative PsychENCODE BW transcriptome score was predictive of BW in the replication samples (GTEx partial r = 0.19, P = 0.0266; ROSMAP partial r = 0.17, P = 1.44e^-5^; **Fig. S12**), demonstrating similar out-of-sample prediction as polygenic scores derived from genome-wide association studies of structural neuroimaging phenotypes in much larger cohorts ^6^. Moreover, we found this BW transcriptome score to have significant group differences across neuropsychiatric disorders (PEC), as well as patients with a reported history of smoking (GTEx; **Fig. S12**) – a well-documented comorbidity with deleterious effects on brain morphology ^7^. Additionally, considering the spatial alignment between the BW +/- relative expression and neurodegenerative disease pathology (from tau PET), we found significant group differences in the BW transcriptome score in patients with dementia (ROSMAP). In ROSMAP, we implemented a linear mixed effects model framework similar to the per gene brain weight models (Methods) with the BW transcriptome score instead as the outcome variable and Braak stage or amyloid as predictor variables in lieu of brain weight. We found significant relationships between interindividual variation in BW transcriptome scores and both postmortem neuropathological metrics (all gene BW transcriptome score: Braak stage t = -2.173, P = 0.0171; amyloid t = -4.832, P = 1.71e^-6^).

#### Lifespan Brain Chart Consortium

Please find a list of members here: <https://github.com/brainchart/Lifespan>.

### Supplemental figures

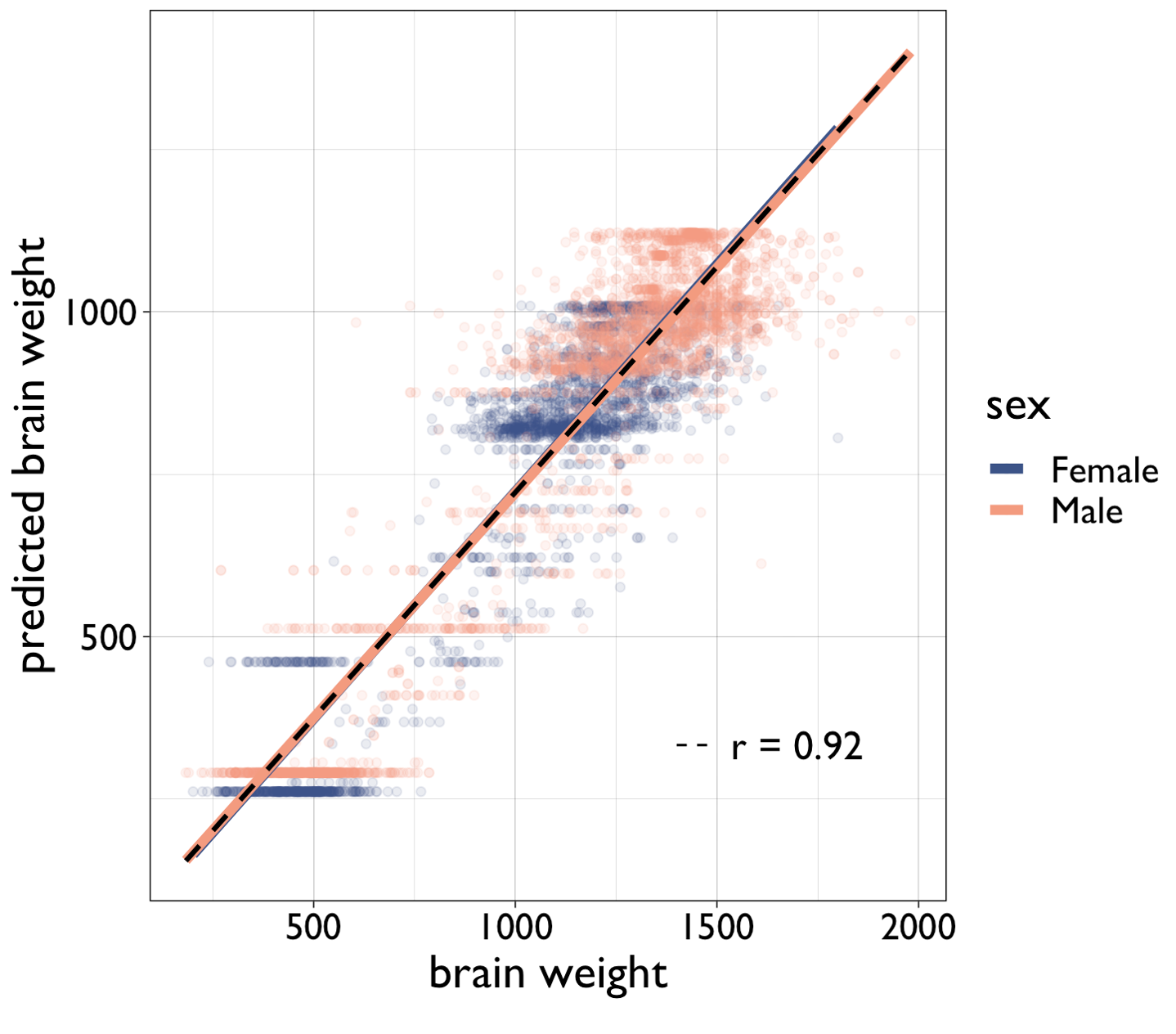

**Figure S1.** Scatterplot showing the relationship between empirical brain weight (grams) and predicted brain weight (grams), based on lifespan neuroimaging models of brain size (total cerebrum volume) from a prior study (Total N=3,689; PsychENCODE N=1,670; GTEx N=227; ROSMAP N=634; independent aggregated dataset without associated transcriptomics N=1,158 ^8^). Colors denote biological sex. Lines show linear fits. Black dotted line shows overall fit (Pearson r=0.92, P < 2.2e-16).

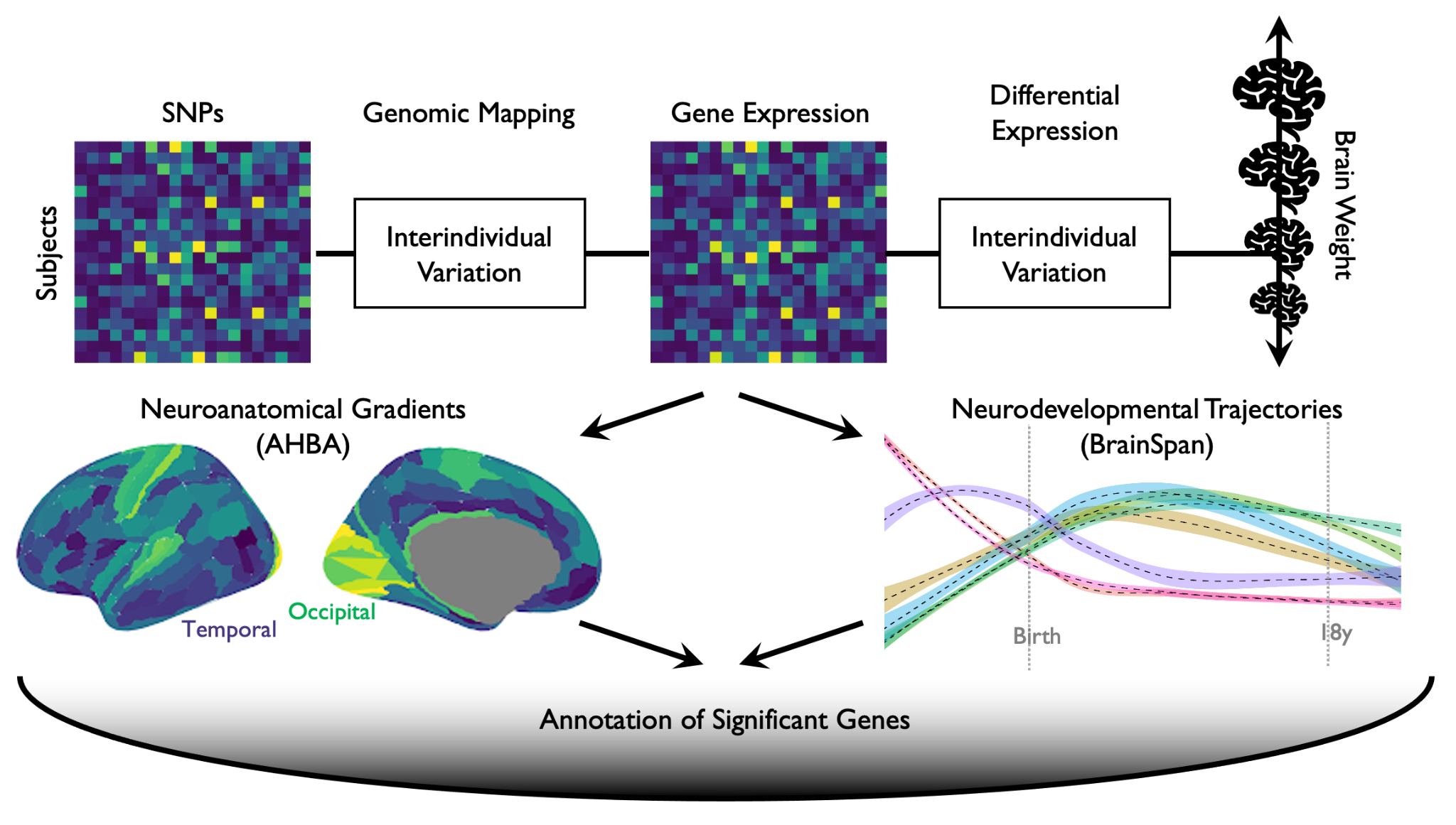

**Figure S2.** Schematic representing the workflow for the current study. SNPs=single nucleotide polymorphisms; AHBA=Allen Human Brain Atlas.

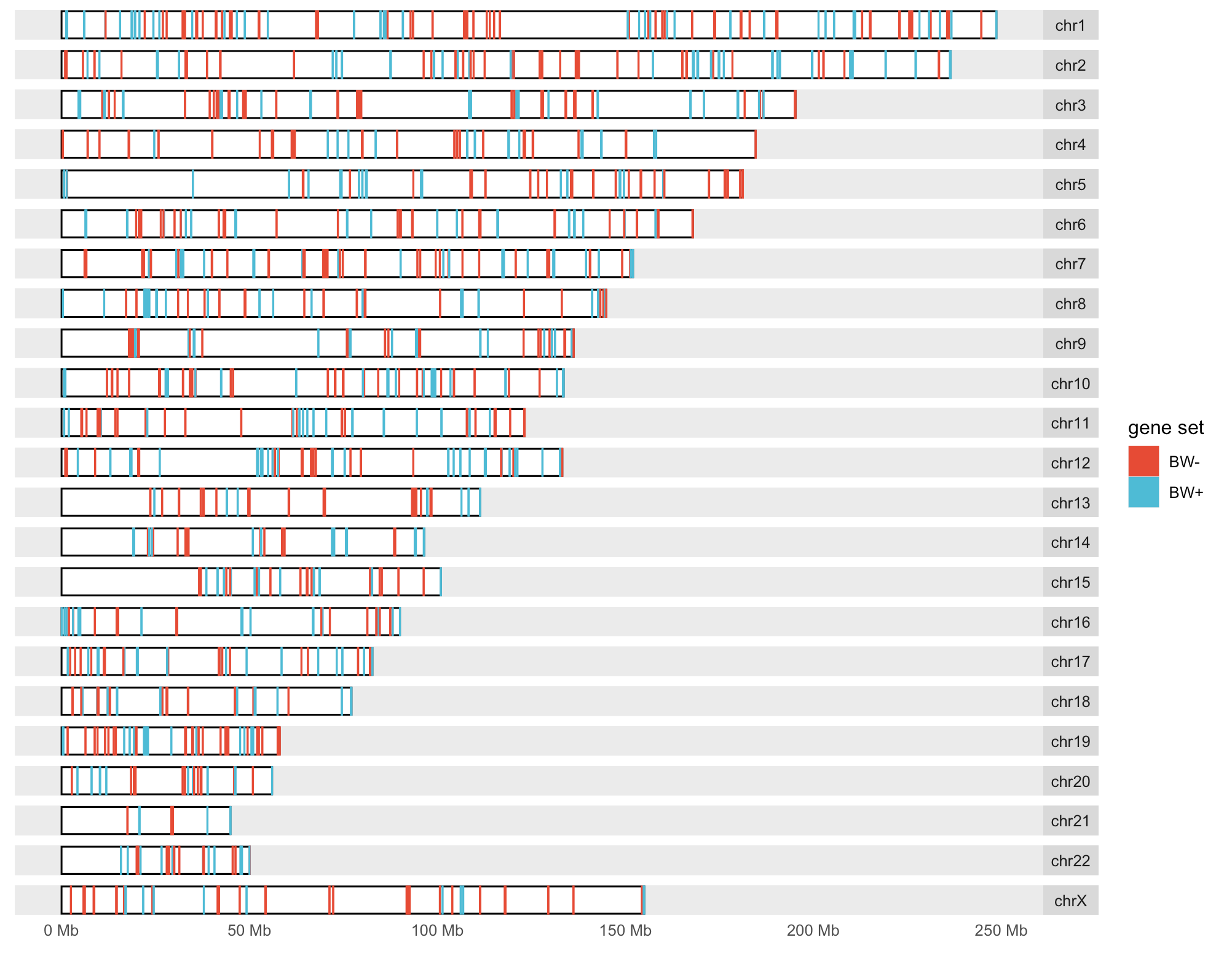

**Figure S3.** Karyogram with colored lines indicating locations of genes with significant (P_Bonferroni_ < 0.05) upregulated expression in smaller-brained (brain weight negative or “BW-”) or larger-brained (brain weight positive or “BW+”) individuals in the PsychENCODE dataset.

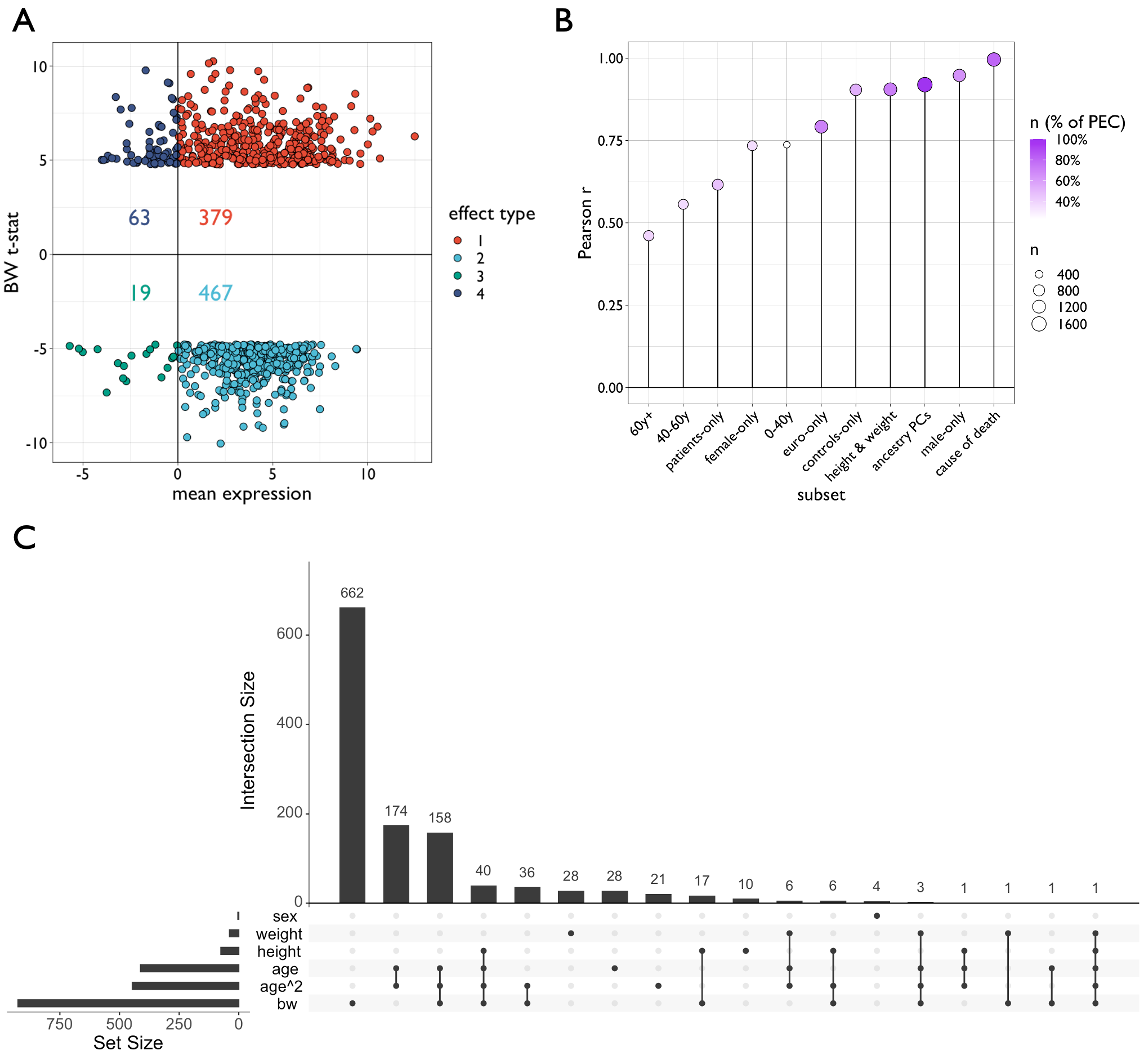

**Figure S4.** Sensitivity analyses of brain weight gene expression models. **A |** scatterplot showing the relationship between the average expression (z-score) and brain weight (BW) association (t-statistic) in the main PsychENCODE (PEC) dataset. Each dot represents a gene that was found to have a significant association (P_Bonferroni_ < 0.05) between brain weight and expression across samples. Colors and numerical types are arbitrary. Both BW+ and BW- genes (positive and negative values on y-axis, respectively) disproportionately reflect genes with generally higher expression (more positive values on x-axis), which allows for the opposing interpretation of the two gene sets. Numbers represent the counts of genes within each type. **B |** Lollipop plot showing the Pearson correlation coefficient across genes between the main PEC model output and output generated based on subsets of the PEC dataset. Circle color is the percentage of the total PEC sample in each subset, and size is the absolute sample size of each subset. 60+y=donors aged 60 years and older, 40-60y=donors aged 40-60 years, patients-only=just donors with a documented psychiatric diagnosis, female-only=just female donors, 0-40y=donors aged 0-40 years, controls-only=just donors without a documented psychiatric diagnosis, height & weight=donors with height and weight information (height and weight included as covariates in models), male-only=just male donors, euro-only=just donors of european ancestry, ancestry PCs=inclusion of ancestry principal component scores based on genotype in gene models, cause of death=cause of death included as a covariate in models. **C |** Upset plot demonstrating the number and overlap of significant (P_Bonferroni_ < 0.05) genes for biological traits in the PsychENCODE discovery cohort. For brain weight (“bw”), sex, linear age (“age”) and squared age (“age^2”) the original model was used. Since only a subset of subjects had documented height and weight, those terms were included as additional covariates in separate gene-level models (e.g., as part of the analyses in panel B).

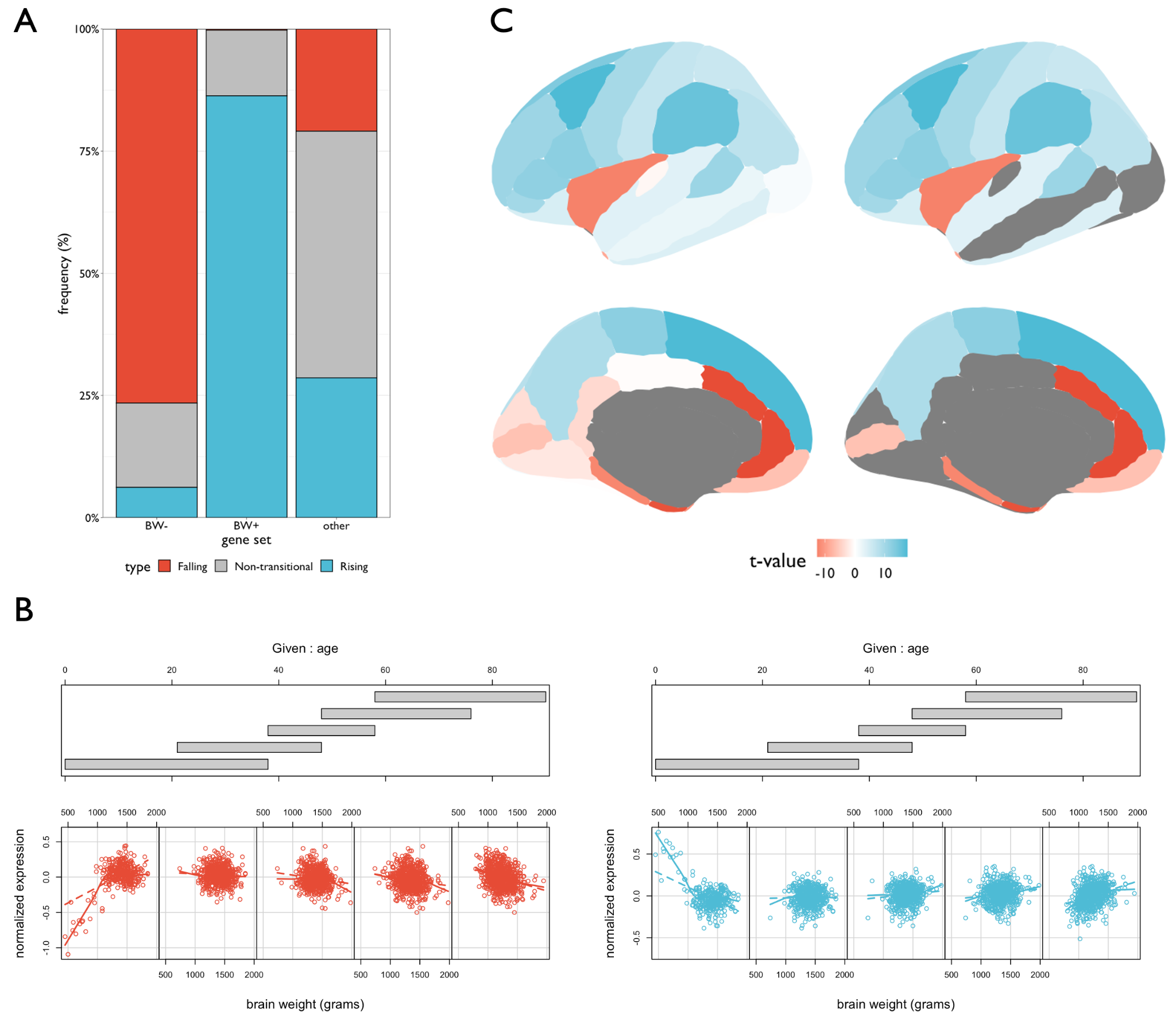

**Figure S5.** Spatiotemporal evaluation of the brain weight associated genes. **A |** Stacked bar plots showing the proportion of genes within each BW gene set that fall within predefined categories of perinatal expression trajectories (Werling et al., 2020). Colors were set based on qualitative assessment of the strong bias for “falling” (BW-, red) and “rising“ (BW+, blue) effects. The “other” gene set type refers to all genes not a part of the BW sets. **B |** Conditioning plots showing the interaction effects of brain weight and age on gene expression. Age ranges (gray bars) were determined based on the distribution of age values. Genes were identified based on showing significant brain weight by age interaction effects (P_Bonferroni_ < 0.05) but not significant age or brain weight effects (both P_Bonferroni_ > 0.05). 92 genes had negative coefficients (red, left) and 130 genes had positive coefficients (blue, right). The number of age strata (N=5) and percent overlap (50%) were arbitrarily chosen for visualization. Dashed lines denote linear fits, while solid lines denote nonlinear (LOESS) fits. **C |** Brain plots showing the BW+/- relative expression differences (t-values), using a coarser cortical parcellation of the Allen Human Brain Atlas. The left column shows the complete set of effects for each region, and the right column shows only significant effects (P_Bonferroni_ < 0.05; gray values P_Bonferroni_ > 0.05). The top row shows the lateral view of the left hemisphere, and the bottom row shows the medial view.

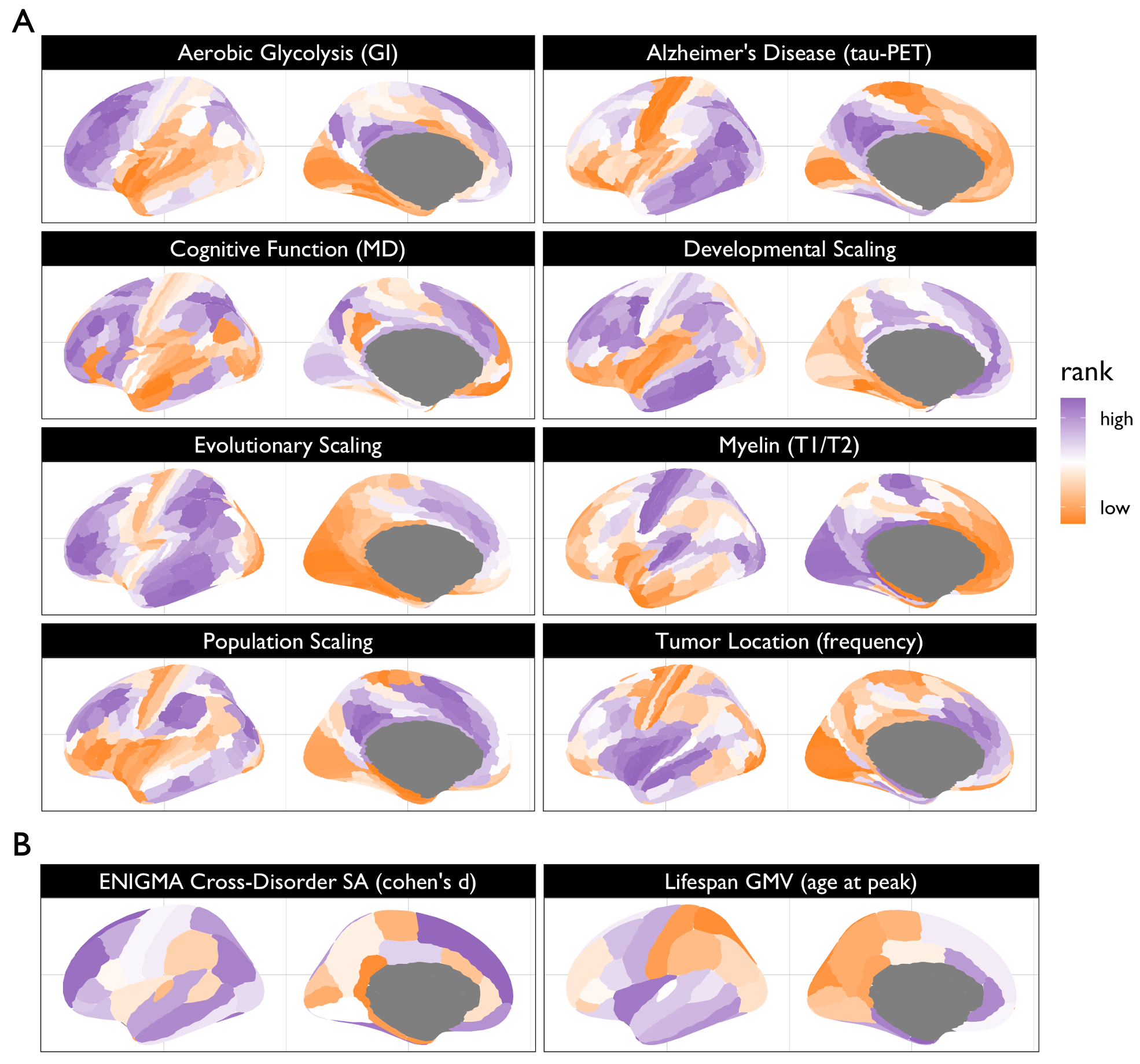

**Figure S6.** **A |** Brain plots showing the cortical region topographies of various neuroimaging maps in a 180-region cortical parcellation ^9^. Each map was bilaterally averaged and plotted on the left hemisphere for visualization. Colors correspond to rank values for each map for visualization purposes (orange=low values, purple=high values). Aerobic glycolysis was measured with the glycolytic index (GI) from PET, Alzheimer’s disease pathology was measured with tau-PET, cognitive function of the multiple demand network (MD) was measured with fMRI, developmental scaling estimated surface area expansion from structural MRI comparing children to adults, evolutionary scaling estimated surface area expansion from structural MRI comparing macaques to humans, myelin was estimated using an established ratio of T1w to T2w structural MRI across subjects, population scaling estimated surface area expansion from structural MRI data across humans, and tumor frequency was calculated using manually drawn masks of glioblastoma and low grade glioma across subjects. **B |** Brain plots plotted similarly as in A, but in a 34-region cortical parcellation. The cross-disorder surface area (SA) map represents the first principal component across available regional summary statistics from the ENIGMA consortium for attention deficit hyperactivity disorder, bipolar disorder, major depressive disorder, obsessive compulsive disorder, and schizophrenia. The lifespan gray matter map represents ages at peak absolute volume (averaged across males and females). See Methods for more information on all the maps.

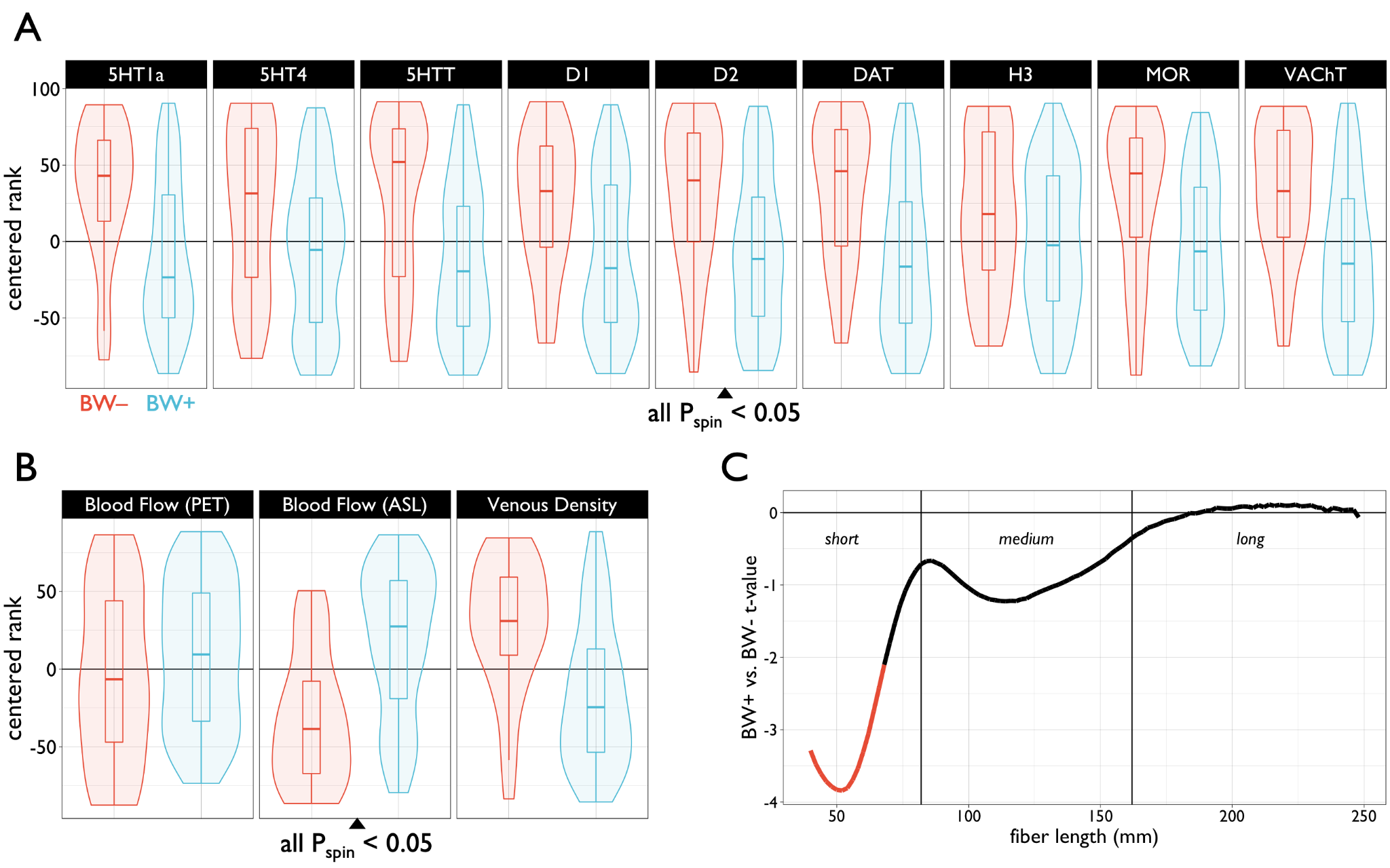

**Figure S7. Heterotopic contextualization of brain weight (BW) differential expression**. **A |** Comparison of independently-measured neurotransmitter densities in significant regions of BW+ (blue) or BW- (red) differential expression in the Allen Human Brain Atlas. Differences between BW+ and BW- were quantified with t-tests, and all comparisons shown were significantly different from statistics generated from spatially-permuted differential expression maps (P_spin_ < 0.05). Note that all relationships show greater BW- values. Centered rank values were computed by ranking (from low to high) regional estimates for each expression type and subtracting the median rank, which was used to aid in comparison across maps. **B |** Same as A, instead comparing areas of differential expression in terms of neuroimaging-derived estimates of cerebral blood flow and venous density. **C |** Comparison of prevalence of white matter fiber lengths across areas of BW differential expression. T-values were computed across BW+ versus BW- regions in terms of fiber density values at varying fiber lengths. The black line denotes fiber lengths where BW regional differences were insignificant (P_Bonferroni_ > 0.05), red where BW- > BW+ (P_Bonferroni_ < 0.05). PET: positron emission tomography, ASL: arterial spin labeling. See Methods for further information on the neurotransmitter maps. All box-violin plots show median and interquartile range (IQR) with whiskers denoting 1.5 × IQR.

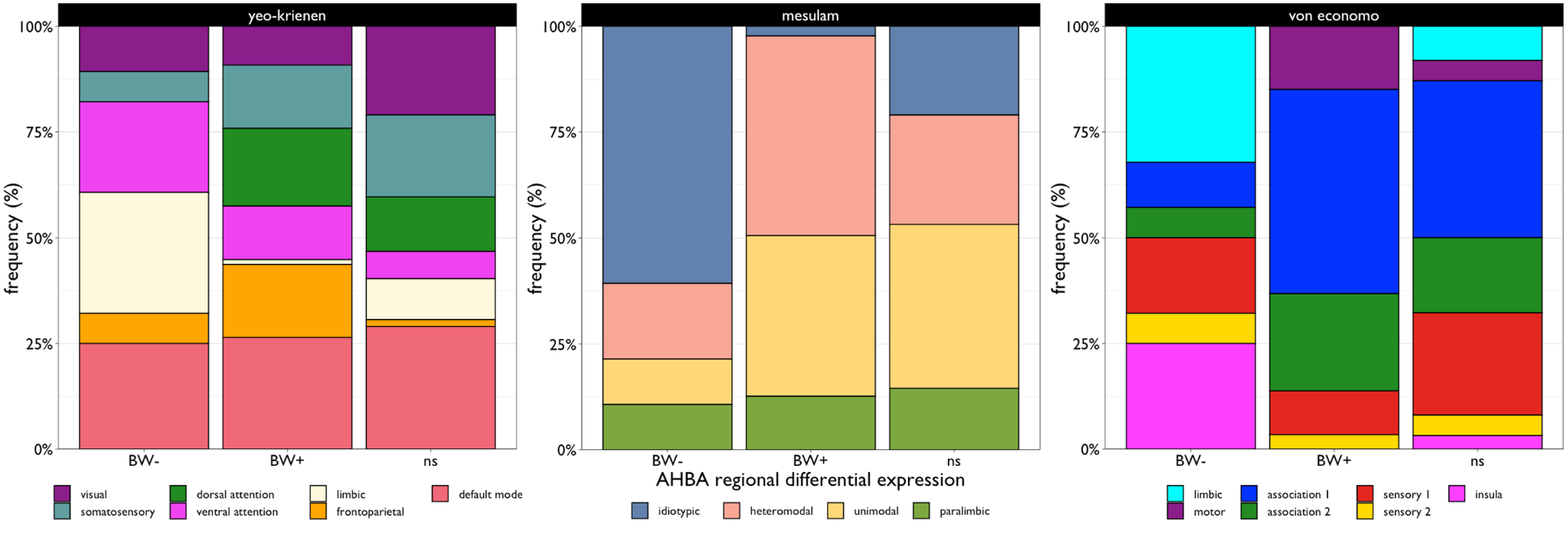

**Figure S8.** Heterotopic categorization of regional brain weight (BW) relative expression (from Fig. 1D). Stacked barplots show the proportion of BW+ and BW- regions based on functional (left), hierarchical (middle), and cytoarchitectonic (right) classification. AHBA: Allen Human Brain Atlas, ns: non-significant regions of differential expression.

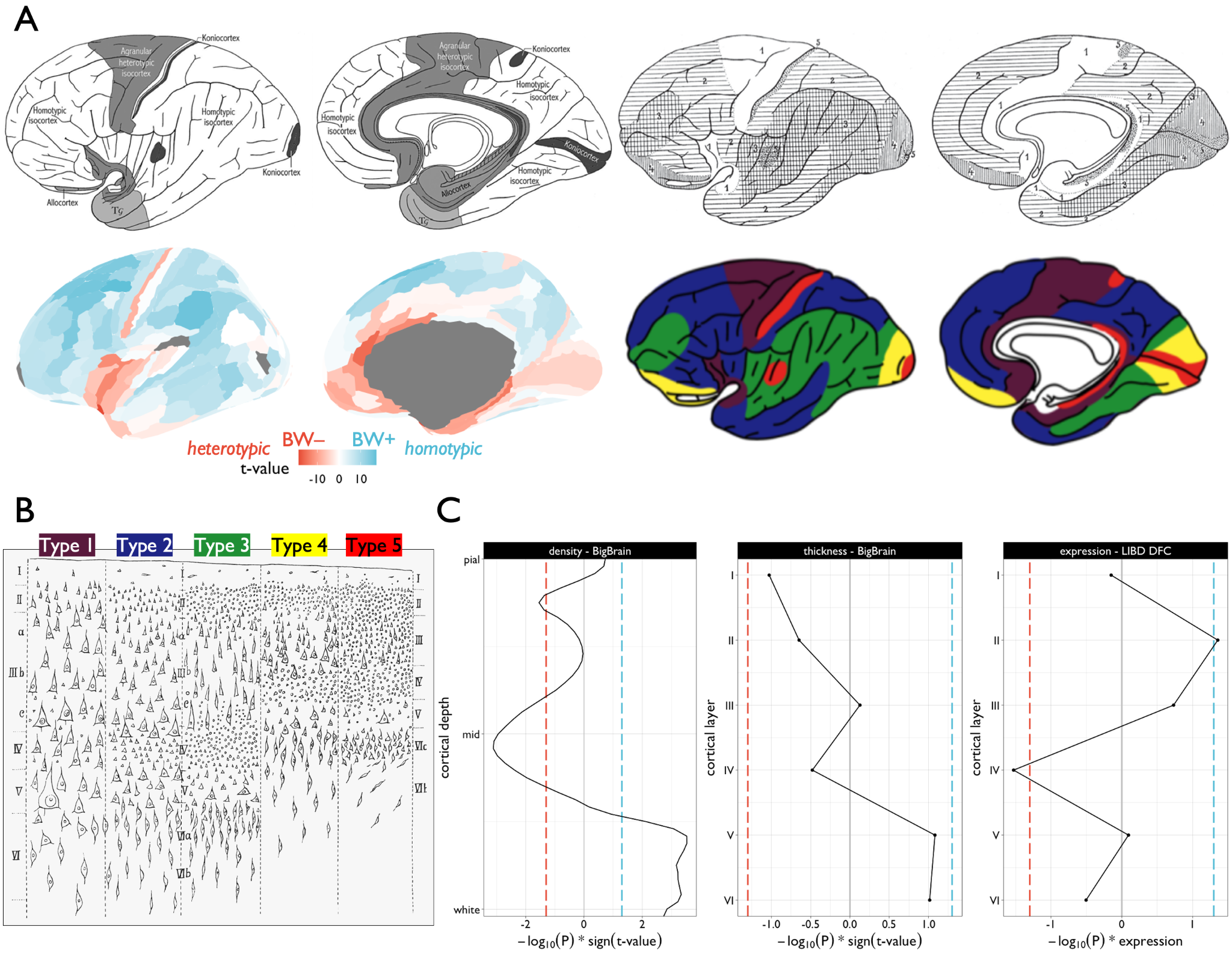

**Figure S9**. **A |** Von Economo-Koskinas cortical atlases ^10^ based on cytoarchitectonic characteristics (top left), and subdivided based on one of five cortical types (top right, colored in bottom right). The full map of the cortical region differential expression (t-statistics) of brain weight positive (BW+) versus brain weight negative (BW-) in the Allen Human Brain Atlas (similar to Figure 1, but unthresholded) is shown for qualitative comparison. **B |** The 5 described von Economo-Koskinas cortical types are shown in terms of their general profiles across layers. Notably, types 1 and 5 – which align with regions of BW- differential expression – are distinct in their agranular vs. granular properties, however both show more indistinguishable layer differentiation. **C |** Laminar profile analysis of BW gene set differential expression in three independent datasets. (Left) laminar density and (middle) thickness based on 3D histological reconstruction in the BigBrain dataset characterizing areas of BW+ versus BW- differential expression (from A, bottom left) across 100 cortical depths (density) and assigned layers (thickness). (Right) differential gene expression of BW+ versus BW- gene sets at each layer from the spatial RNA sequencing postmortem brain dataset (in the dorsolateral prefrontal cortex, DFC) from the Lieber Institute on Brain Development (LIBD). Colored dashed lines denote the negative log-scaled p-value significance thresholds (uncorrected P < 0.05, signed by the direction of effect) for BW- (red, negative) and BW+ (blue, positive) relative expression differences. See Methods for more information on these datasets.

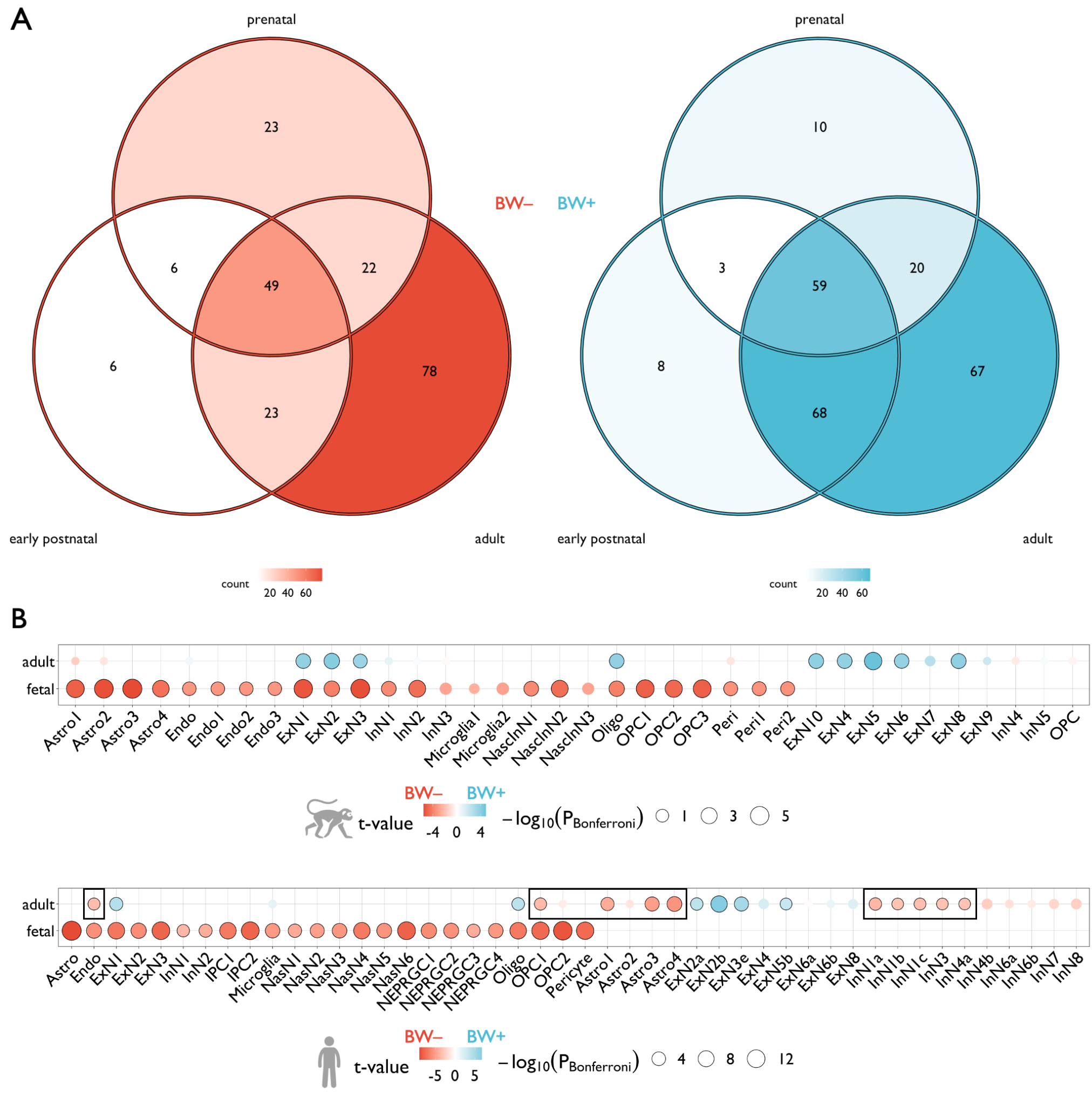

**Figure S10.** **A |** Venn diagrams of brain weight negative (BW-) and brain weight positive (BW+) gene sets based on the significantly differentially expressed genes (P_Bonferroni_ < 0.05) in humans relative to macaques in three developmental epochs. **B |** Relative expression of BW+ versus BW- genes across individual cell-types, using cell-specific RNA sequencing data in fetal and adult samples from macaques (top) and humans (bottom). BW- relative expression (red) indicates that BW- genes are more highly expressed in that cell type compared to BW+, whereas BW+ relative expression (blue) indicates the opposite effect. Black outlines denote significant effects (P_Bonferroni_ < 0.05). Circles are scaled according to Bonferroni-corrected p-values. Black rectangles denote human-specific effects relative to macaques. X-axis labels are arbitrarily ordered based on alphabetized fetal cell subtype labels. Alignment of adult cell subtype labels is purely based on shared naming, for visualization purposes. Astro: astrocytes, Endo: endothelial cells, ExN: excitatory neurons, InN: inhibitory neurons, NasN/NascInN: nascent neurons, Oligo: oligodendrocytes, OPC: oligodendrocyte precursor cells. Peri: pericytes.

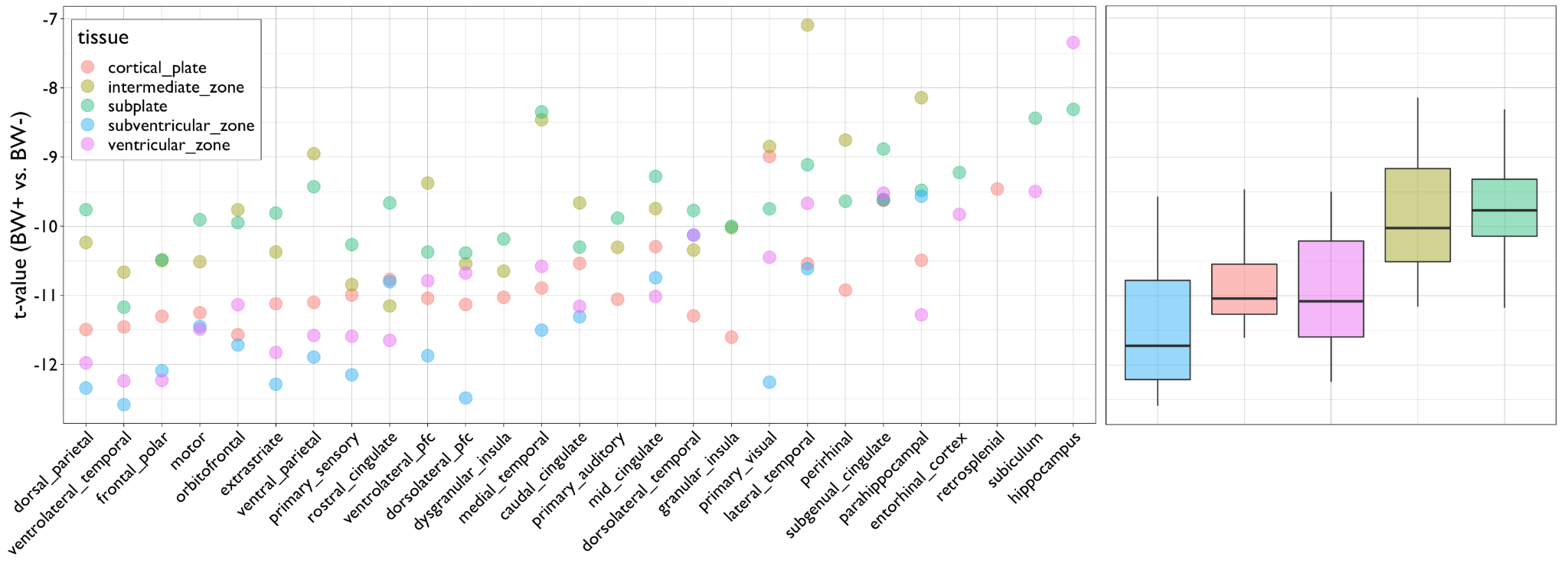

**Figure S11.** Differential expression of brain weight positive (BW+) versus brain weight negative (BW-) gene sets in an independent prenatal developmental dataset (BrainSpan) across brain areas and zones during midgestation. T-statistics show all areas (circles, left) and zones (boxplots, right) show significant BW- differential expression, with notable gradated variation across zones – the subventricular zone showing greatest BW- differential expression relative to the other tissue types (t = -2.22, P = 0.029). All boxplots show median and interquartile range (IQR) with whiskers denoting 1.5 × IQR.

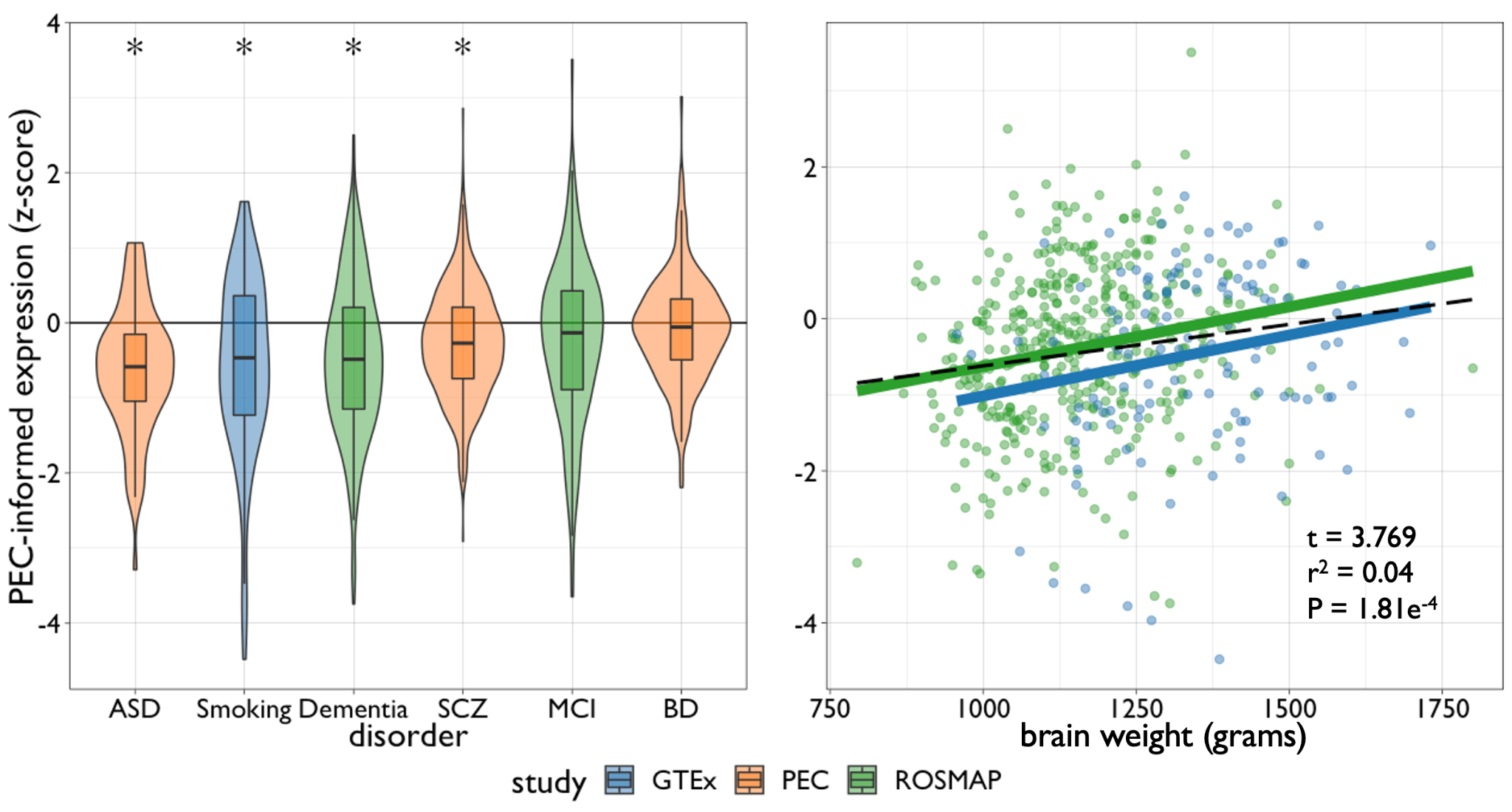
**Figure S12**. (Left) distribution of brain weight (BW) transcriptome scores in patients with a diagnosed psychiatric or neurological disorder. BW transcriptome scores were computed using the PsychENCODE (PEC) BW model coefficients for all genes. Scores for each subject were generated by summing the weighted expression across all genes (dot product of all gene-wise coefficients of the PEC brain weight models and gene expression), and normalized (z-scored) relative to their respective study controls. Smoking refers to smoking status, recorded as a binary variable upon autopsy. ASD: Autism Spectrum Disorder, AD: Alzheimer Disease, SCZ: Schizophrenia, MCI: Mild Cognitive Impairment, BD: Bipolar Disorder. Asterisks represent nominal significance (uncorrected P < 0.05) of disorder groupwise mean score compared to zero. All box-violin plots show median and interquartile range (IQR) with whiskers denoting 1.5 × IQR. (Right) Scatterplot of the relationship between BW and BW transcriptome score. Black line and reported statistics represent the fit across datasets using a linear mixed effect model with fixed effects of age and sex, and random effects of diagnostic category (see left panel) and study.

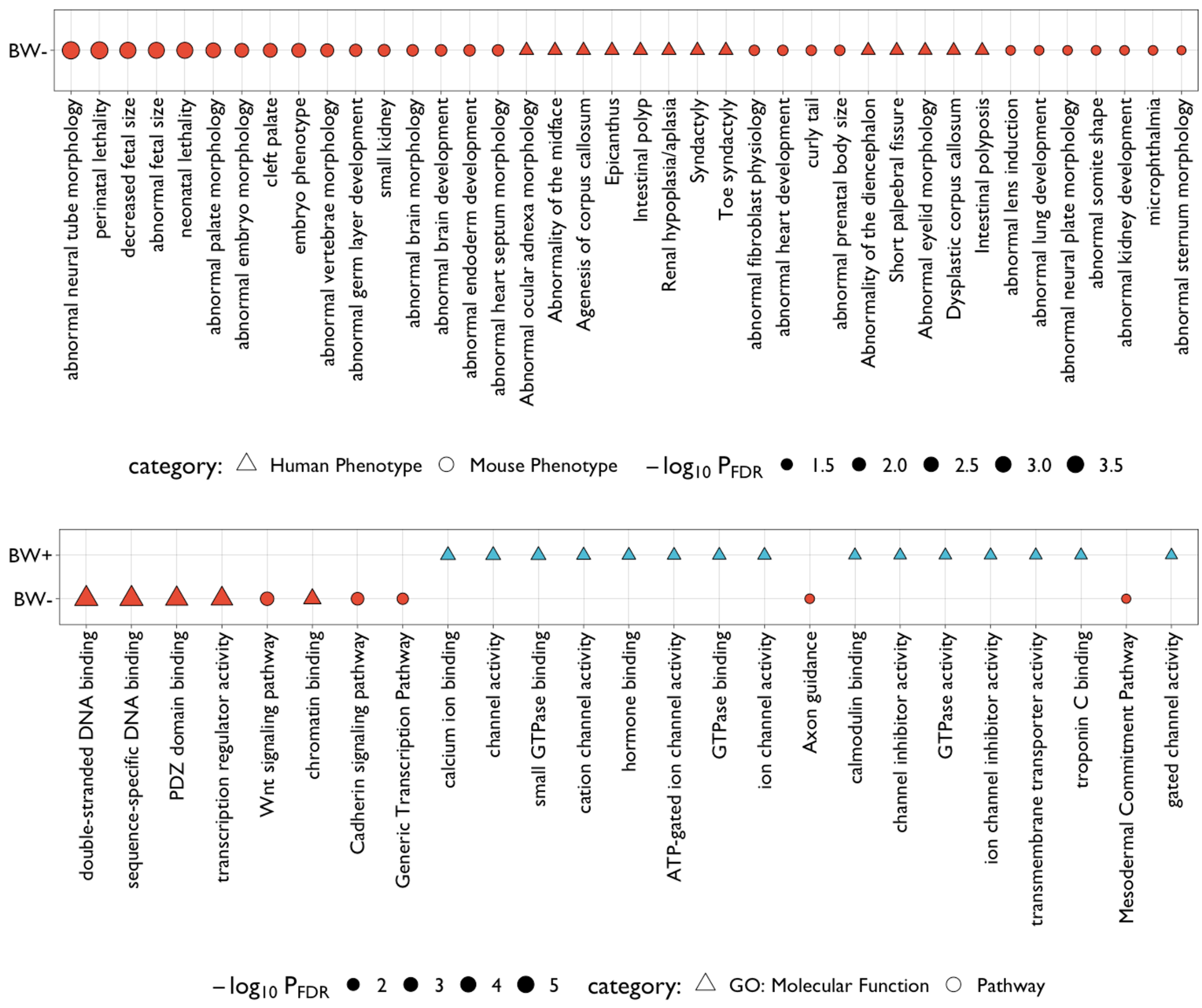

**Figure S13**. Grid plot showing significant (P_FDR_ < 0.05, see Table S4) gene set enrichment of brain weight (BW) associated genes for Human and Mouse Phenotypes (top) and GO: Molecular Function and Pathway (bottom) using ToppGene. Shapes are sized according to adjusted negative log-scaled p-values.

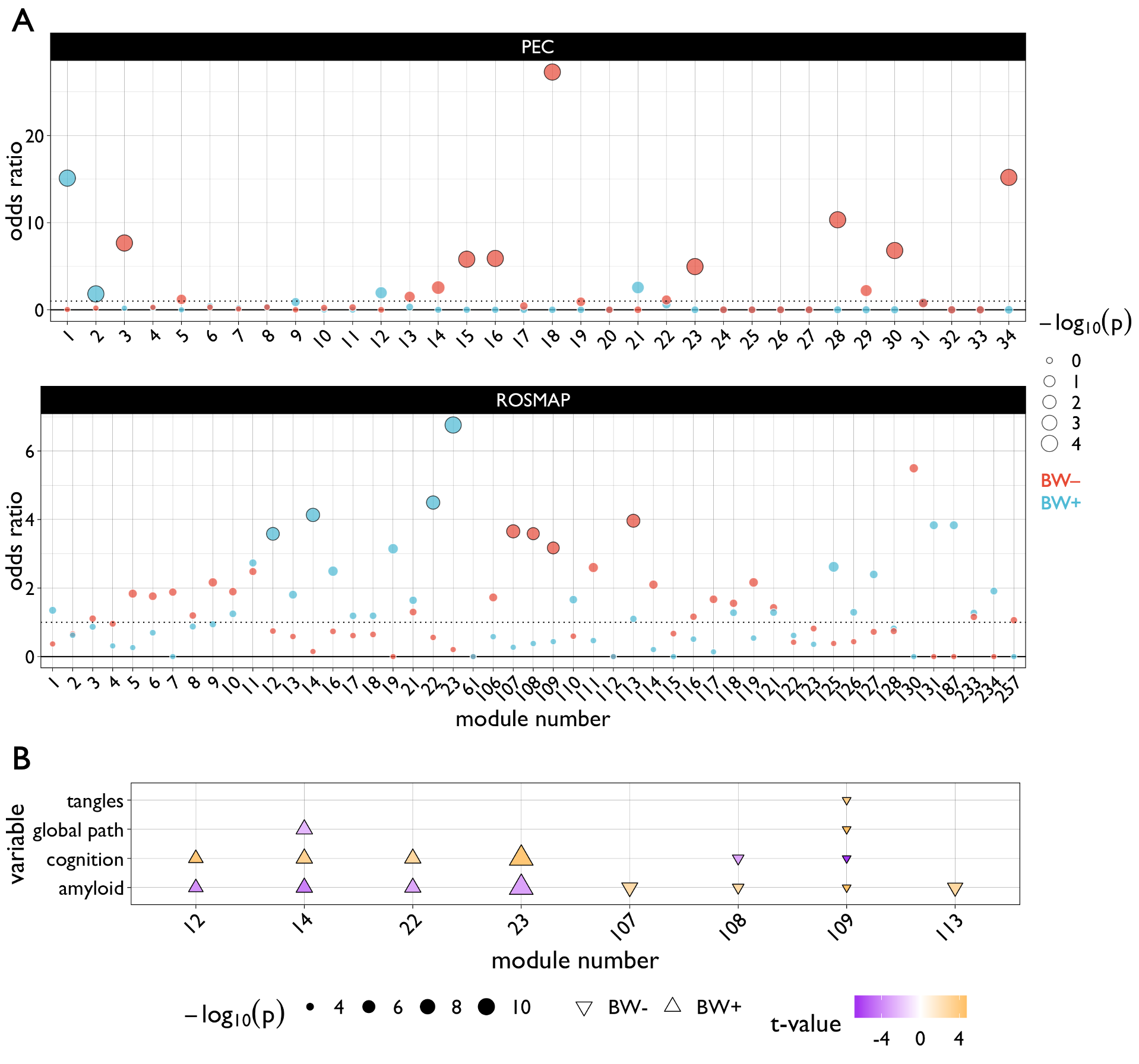

**Figure S14. A |** Enrichment of brain weight (BW) gene sets in gene co-expression modules in the PsychENCODE (PEC) and ROSMAP datasets. Modules were derived in previous work ^11,12^, and gene set enrichment was calculated using hypergeometric tests. The size of the circles denotes negative log-scaled uncorrected p-values, with color corresponding to the BW gene set. Circles with black outlines are significant at P_FDR_ < 0.05 (corrected across number of modules). Dashed line denotes odds ratio = 1. See also Table S5 from the original PEC paper ^11^ which generated these module classifications for cell-type and disease enrichments. **B |** Annotations of significant (P_FDR_ < 0.05) BW-enriched modules based on metrics derived from postmortem neuropathology in the ROSMAP cohort. Shapes correspond to which BW gene set is enriched in a given module, and colors represent t-statistics from linear models comparing module eigengene scores to each neuropathology metric across individuals. Only uncorrected P < 0.05 effects are shown for visualization. See Table S6 in this paper for complete statistics in the ROSMAP cohort.

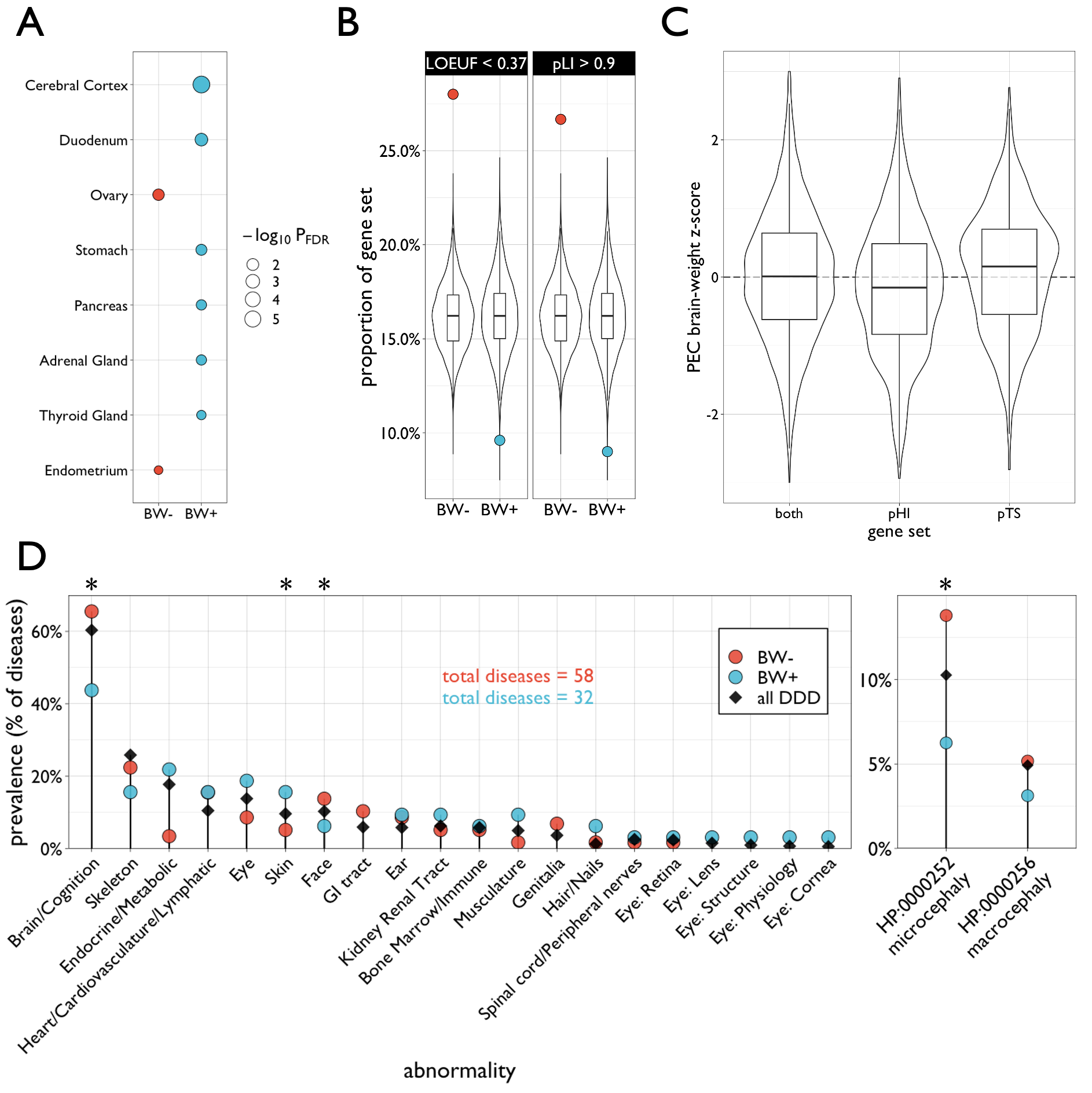

**Figure S15. A |** Gene set enrichment analysis (hypergeometric test) of brain weight (BW) gene sets across tissues in the Human Protein Atlas. For visualization only significant (P_FDR_ < 0.05) results are shown. Circles are colored according to BW gene sets and sized according to negative log-scaled p-values. **B |** Enrichment analysis of BW gene sets based on proportion of genes showing high loss-of-function intolerance based on two validated metrics derived from an external whole exome sequencing dataset (gnomAD v2.1.1). Recommended thresholds of each metric allowed for the LOEUF and pLI scores to be comparable. Colored circles represent the proportion of genes in BW gene sets meeting each established threshold (LOEUF < 0.37 or pLI > 0.9), and box-violin plots show the proportion overlap distribution across resampled 10,000 gene sets of similar size to the BW+ and BW- sets. **C |** Box-violin plots showing differential gene expression coefficients for the brain weight analysis in PsychENCODE (PEC) across gene sets based on dosage sensitivity probabilities for haploinsufficiency (pHI) and triplosensitivity (pTS) . Gene sets were established using predefined dosage sensitivity thresholds (pHI > 0.86 and pTS > 0.94). The pHI and pTS gene sets were further refined to exclude genes meeting both probability thresholds (constituting the “both” category). **D |** Prevalence of associated phenotypic symptomatology in diseases with de novo mutations in overlapping BW genes in the Deciphering Developmental Disorders (DDD) dataset (DECIPHER v11). Colored circles represent the prevalence of respective phenotypes within the 58 developmental disorders of overlapping BW- genes and 32 developmental disorders of overlapping BW+ genes. Black diamond represents the prevalence of respective phenotypes across all documented disorders in the database. Cephalic conditions were separated out based on identification via associated Human Phenotype (HP) documentation. Red and/or blue circles may be obscured for some low prevalence phenotypes (e.g., “Eye: Lens”). Asterisks denote significant enrichment for a BW gene set above expected based on prevalence across all disorders (black diamond) and permutation testing (P_permutation_ < 0.05). All box-violin plots show median and interquartile range (IQR) with whiskers denoting 1.5 × IQR. See Methods for more information on these datasets.

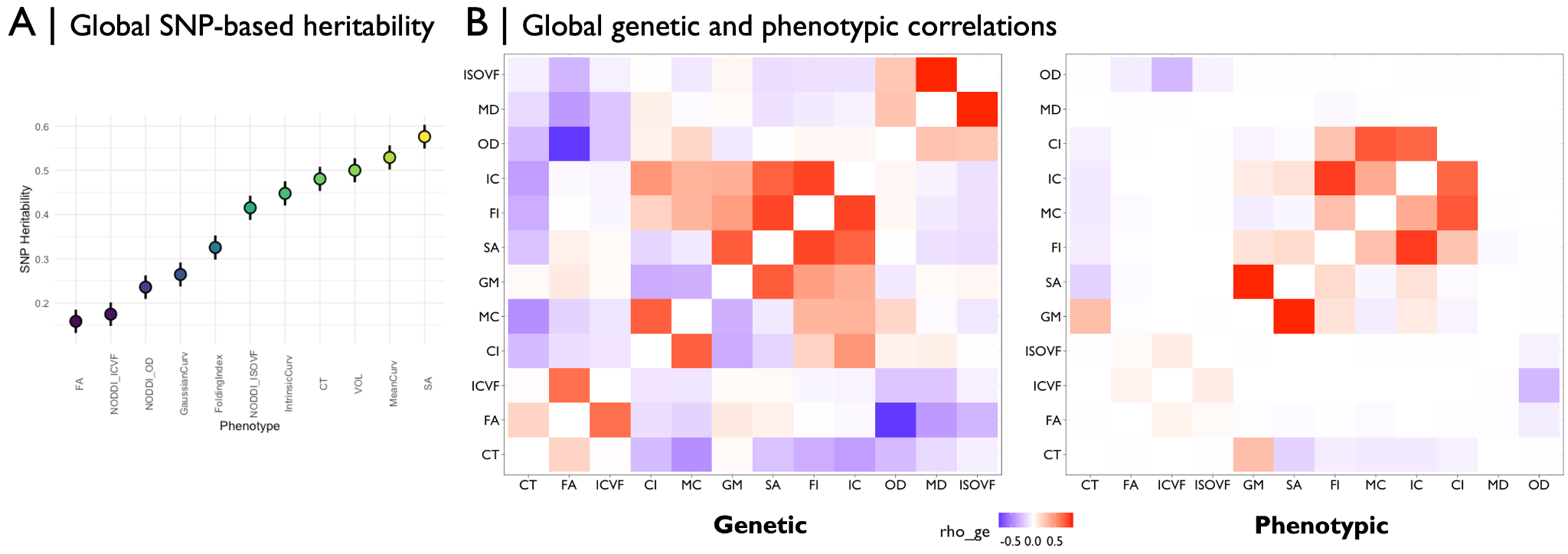

**Figure S16. A |** SNP-based heritability of global multimodal neuroimaging phenotypes (averaged or summed across the brain) in the UK Biobank. **B |** Genetic (left) or phenotypic (right) correlations (“rho_ge”, using the ‘RHOGE’ package; <https://github.com/bogdanlab/RHOGE>) across the transcriptome-wide association study summary statistics for each global neuroimaging phenotype. FA=fractional anisotropy, NODDI_ICVF=intracellular volume fraction derived from neurite orientation dispersion and density imaging (NODDI), NODDI_OD=orientation dispersion derived from NODDI, GaussianCurv=Gaussian curvature, IntrinsicCurv=intrinsic curvature, NODDI_ISOVF=isotropic volume fraction derived from NODDI, CT=cortical thickness, VOL=gray matter volume, SA=surface area.
